# CLPC2 plays specific roles in CLP complex-mediated regulation of growth, photosynthesis, embryogenesis and response to growth-promoting microbial compounds

**DOI:** 10.1101/2025.11.25.690394

**Authors:** Jesús Leal-López, Abdellatif Bahaji, Nuria De Diego, Petr Tarkowski, Edurne Baroja-Fernández, Francisco José Muñoz, Goizeder Almagro, Carlos Eduardo Aucique-Pérez, Francisco Ignacio Jasso-Robles, Domenico Loperfido, Elisabetta Caporalli, Ignacio Ezquer, Lidia López-Serrano, Alberto Férez-Gómez, Víctor Coca-Ruiz, Pablo Pulido, Rafael J. L. Morcillo, Javier Pozueta-Romero

## Abstract

In Arabidopsis, exposure to growth-promoting microbial volatile compounds (VCs) enhances CLPC2 levels. This chaperone forms part of the CLP protease complex, which ensures the correct functioning of essential processes in plastids. Previous studies indicated considerable functional redundancy of CLPC2 with its dominant paralogue CLPC1. However, the function and action mechanism of CLPC2 still remain unknown. Here we found that CLPC2-lacking *clpc2-2* mutants were unresponsive to microbial VCs, whereas *clpc1-1* knockout mutants exhibited a WT-like response to VCs when grown on sucrose-containing medium. Unlike *clpc1-1*, *clpc2-2* plants presented a fully functional photosystem II and lower than WT stomatal conductance. Furthermore, *clpc2-2* plants, but not *clpc1-1* plants, produced wrinkled seeds with delayed embryonic development and reduced postgerminative establishment rates that resembled those of mutants lacking P and R components of the CLP proteolytic core. Proteomic analyses revealed that knocking out of *CLPC2* enhanced the levels of chloroplastic proteins that are essential for growth, embryo development and seedling establishment. These changes differed from those promoted by the lack of CLPC1, but partially resembled those promoted by CLPPR core inactivation. Nearly 40% of the proteins differentially accumulated by the lack of CLPC2 were VC-responsive. Notably, *35S* promoter-driven *CLPC2* expression promoted changes in the proteome similar to those promoted by the lack of CLPC1. Collectively, our findings highlighted contrasting functional and molecular specificities for CLPC1 and CLPC2, and provided strong evidence that CLPC2 plays specific roles in CLP complex-mediated regulation of plant growth, photosynthesis, embryogenesis, postgerminative seedling establishment and microbial VC responsiveness.

## INTRODUCTION

The lifespan and activity of proteins depend on protein quality control (PQC) systems formed by chaperones and proteases that ensure correct protein folding and prevent the formation of toxic aggregates. These systems operate throughout the growth and development of the plant, and during adaptation to stress insults in which unfolded polypeptides might aggregate and disrupt the correct functioning of essential metabolic pathways (Isono et al., 2024). In chloroplasts, PQC is mainly mediated by three classes of ATP-dependent proteases that also have chaperone activities: long filament phenotype (Lon), filamentation temperature-sensitive H (FtsH) and caseinolytic protease (CLP) (Nishimura and van Vijk, 2015). In Arabidopsis, CLP is a hetero-oligomeric complex composed of several functional units, including a tetradecameric protease CLPPR core comprising proteolytically active (CLPP1 and CLPP3-6) and inactive (CLPR1-4) subunits; accessory proteins (CLPT1-2) for assembly and stabilization of the CLP core; a hexameric regulatory complex of chaperones with ATPase and CLP core binding domains (CLPC1, CLPC2 and CLPD) that participates in the recognition, unfolding, and translocation of the substrate to the catalytic chamber for degradation; and adaptors or recognins (CLPS1 and CLPF) that modulate the recognition and delivery of the substrate to the chaperone core (Nishimura and van Vijk, 2015). Components of the CLPPR core play essential functions in embryogenesis, plastid biogenesis, and plant development (Kim et al., 2009; Zybailov et al., 2009; Kim et al., 2013), whereas CLPC1 is required for housekeeping processes such as chloroplast development, chlorophyll and carotenoid biosynthesis, and protein import (Constan et al., 2004; Sjögren et al., 2004; Kovacheva et al., 2007; Flores-Pérez et al., 2016; Welsch et al., 2018). Consistently, CLPC1 inactivation causes stunted growth, leaf chlorosis and reduced photosynthesis (Constan et al., 2004; Sjögren et al., 2004; López et al., 2024). Proteomic characterization of *clpc1* mutants and *in vivo* trapping of proteins interacting with CLPC1 studies revealed that the interactome of CLPC1 includes all members of the CLP system, enzymes of the methylerythritol phosphate (MEP) and shikimate pathways and proteins involved in chloroplast proteostasis, chlorophyll biosynthesis, and DNA/RNA, fatty acid (FA), thiamine-PP and sulfur metabolisms (Nishimura et al., 2013; Liao et al., 2022).

The function and action mechanisms of CLPC2 are less understood than those of CLPC1. Park and Rodermel (2004) showed that CLPC2 is a suppressor of the requirement for FtsH in thylakoid biogenesis and photosystem II maintenance during photoinhibition. Some studies showed that *clpc2* knockout mutants display an external phenotype similar to that of wild-type (WT) plants (Constan et al., 2004; Kovacheva et al., 2007; Welsch et al., 2018). This, together with the fact that (i) the *clpc1clpc2* double knockout genotype, but not the single *clpc1* and *clpc2* knockout genotypes, causes embryo lethality in Arabidopsis (Kovacheva et al., 2007), and (ii) CaMV 35S promoter-driven *CLPC2* overexpression can complement the *clpc1* mutation (Kovacheva et al., 2007) indicated a considerable functional redundancy of CLPC1 and CLPC2, although a greater prominence was attributed to CLPC1 as it is expressed more strongly than CLPC2 (Constan et al., 2004; Kovacheva et al., 2007). However, recent studies showed that CLPC2-lacking *clpc2-2* plants have a semi-dwarf phenotype, and provided evidence that CLPC1 and CLPC2 have evolved some specific roles, despite sharing 91% amino acid sequence identity (López et al., 2024).

Microorganisms emit volatile compounds (VCs) that act as efficient semiochemicals in interkingdom communication (Gámez-Arcas et al., 2022a). Recently, we found that some of these VCs (e.g. ethylene, NO, CO and SH_2_) act as signaling molecules that promote plant growth and developmental changes, photosynthesis and metabolic changes including accumulation of MEP pathway-derived cytokinins [CKs] (Sánchez-López et al., 2016a,b; García-Gómez et al., 2019; Gámez-Arcas et al., 2022a; Morcillo et al., 2024). In Arabidopsis, fungal VC exposure triggers extensive transcriptome, proteome and redox-proteome changes that are mediated by chloroplast-to-nucleus retrograde signaling mechanisms (Sánchez-López et al., 2016a,b; Ameztoy et al., 2019; Ameztoy et al., 2021; García-Gómez et al., 2020; Gámez-Arcas et al., 2022a; Gámez-Arcas et al., 2025). A striking alteration in the proteome of leaves of plants exposed to fungal VCs involves the accumulation of CLP client proteins (Ameztoy et al., 2021). VC exposure also reduced the levels of several plastidial FtsH and increased those of chaperonins and co-chaperonins that are involved in the folding and guidance of newly imported plastid proteins into their native states. This indicated that the response of the chloroplast proteome to fungal VCs is mediated by dynamic balances between the expression of chaperones, the CLP protease complex and other proteases. In support of this hypothesis, a *clpr1-2* knockdown mutant of the CLPPR core poorly responded to fungal VCs (Ameztoy et al., 2021). Notably, fungal VC exposure strongly enhanced CLPC2 levels (Ameztoy et al., 2021). This suggested that CLPC2 could play important and specific roles in the adaptation of plants to fungal VCs, as well as in VC-sensitive parameters such as growth, development, photosynthesis, and metabolism. To test this hypothesis and explore possible functional specifities of CLPC1 and CLPC2, here we characterized the responses of *clpc2-2* and *clpc1-1* knockout mutants to fungal VCs. We also compared growth, photosynthesis and seed development in these mutants. Furthermore, to understand the molecular action mechanisms of CLPC2, we compared the proteomic phenotype of *clpc2-2* with those of WT, *clpc1-1* and *clppr* plants. Collectively, our findings highlighted contrasting functional specificities and molecular mechanisms of action for CLPC1 and CLPC2, and show that CLPC2 regulates plant growth, photosynthesis, embryogenesis and response to microbial signals through CLP protease complex action.

## RESULTS AND DISCUSSION

### CLPC2 is required for the fungal VC-promoted growth response

Whether CLPC2 mediates the response of plants to fungal VCs was investigated by characterizing the growth response of *clpc2-2* plants grown on solid MS medium with or without sucrose supplementation to VCs emitted by the fungus *Alternaria alternata*. We also characterized *clpc1-1* plants and *clpc2-2* plants expressing *CLPC2* under the control of its own promoter (*clpc2-2 promCLPC2:CLPC2*). Towards this end, we used the “box-in-box” co-cultivation system in which plants are grown in the vicinity of microbial cultures (Gámez-Arcas et al., 2022a). In the absence of fungal VCs, *clpc1-1* and *clpc2-2* plants grown on sucrose-lacking medium were slightly smaller than WT and *clpc2-2 promCLPC2:CLPC2* plants (**Figure 1**). Sucrose supplementation reverted the semi-dwarf growth phenotype of *clpc1-1* plants back to that of WT (**Figure 1**), strongly indicating that growth retardation of *clpc1-1* plants is mainly due to impairments in photosynthate production. In contrast, this treatment negatively affected growth of *clpc2-2* plants (**Figure 1**). In keeping with Ameztoy et al. (2021), *clpc1-1* plants weakly responded to fungal VCs when grown on sucrose-lacking MS medium. However, these plants exhibited a WT-like growth response to fungal VCs when grown on sucrose-containing medium (**Figure 1**), strongly indicating that CLPC1 predominantly mediates growth responses to fungal VC through mechanisms involving photosynthate production. Regardless of the inclusion of sucrose in the culture medium, *clpc2-2* plants did not respond to fungal VCs (**Figure 1**). Therefore, CLPC2 is an important mediator of plant responses to fungal VCs. The fact that the growth responses of *clpc1-1* and *clpc2-2* to exogenous sucrose and fungal VCs are different strongly indicated functional specificities between CLPC1 and CLPC2.

**Figure 1:**
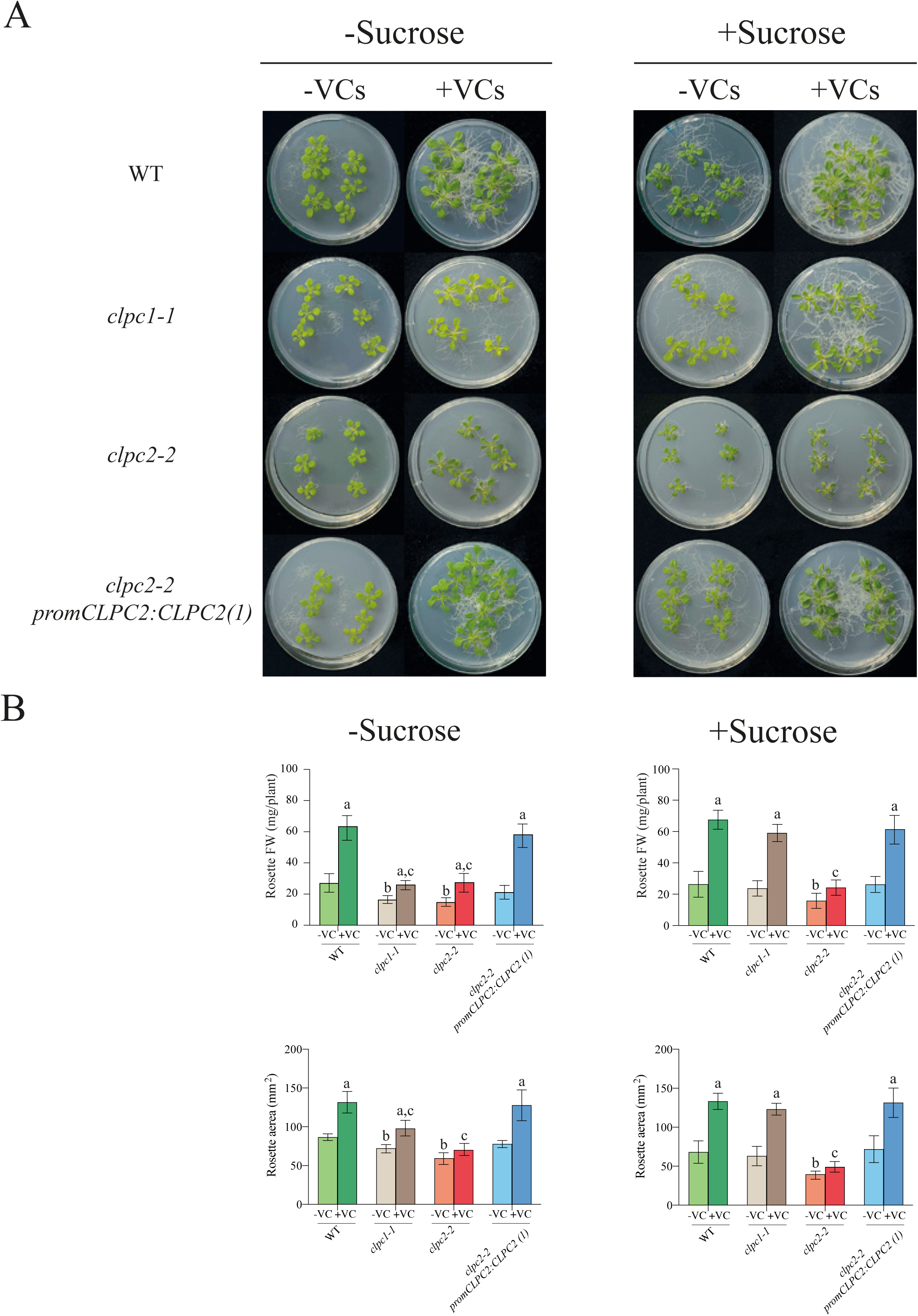
Fungal VCs do not promote growth of *clpc2-2* plants. (A) External phenotype and (B) rosette fresh weight (FW) and area of WT, *clpc1-1, clpc2-2* and one representative line (line 1) of *clpc2-2 promCLPC2:CLPC2* plants grown on solid half-strength MS medium with or without sucrose supplementation in the absence or continuous presence of volatile compounds (VCs) released by adjacent *A. alternata* cultures for one week. Values in “B” are means ± SE for three biological replicates (each comprising a pool of 12 plants) from three independent experiments. Letters “a”, “b” and “c” indicate significant differences, according to Student’s t-test (P<0.05), between: “a” VC-treated and non-treated plants, “b” WT and mutant plants grown without fungal VC treatment, and “c” VC-treated WT and VC-treated mutant plants.

### CLPC1 and CLPC2 mediate photosynthesis through different mechanisms

We carried out time-course analyses of fresh weight (FW) of rosettes of WT, *clpc1-1, clpc2-2* and *clpc2-2 promCLPC2:CLPC2* plants grown on soil. Like *clpc1-1, clpc2-2* plants exhibited a slow growth phenotype, but the WT growth rates could be restored by *CLPC2* expression under the control of the *CLPC2* promoter (**Supplemental Figure 1**). The low growth rates of *clpc1-1* and *clpc2-2* plants suggested the possible occurrence of reduced photosynthetic capacity. We thus measured key parameters of photosynthesis in leaves of WT, *clpc1-1, clpc2-2* and *clpc2-2 promCLPC2:CLPC2* plants grown on soil. Regarding the fluorescence-related parameters, the maximum quantum yields of the photosystem II (PSII) in the dark-adapted state (*F*_v_/*F*_m_) and the operating efficiency of the PSII (<λ_PSII_) values in *clpc1-1* plants were significantly lower than those of WT plants (**Figure 2A,B**). Furthermore, <λ_NPQ_ values in *clpc1-1* plants were higher than those of WT plants (**Figure 2A,B**). This indicated that *clpc1-1* leaves used the light less efficiently, had a more deficient electron transport downstream from PSII and, therefore, produced less NADP^+^ for CO_2_ assimilation than WT leaves. Consistently, *A_n_* values of *clpc1-1* leaves were 40-50% lower than those of WT leaves (**Figure 2C**). Furthermore, *g_s_* and *E* values in *clpc1-1* plants were higher than those of WT plants (**Figure 2C**). Consequently, the values of intrinsic and instantaneous water use efficiencies (WUE-int and WUE-ins, respectively) of *clpc1-1* leaves were higher than those of WT plants. Unlike in *clpc1-1* plants, *Fv/Fm*, <λ_PSII_ and <λ_NPQ_ values in *clpc2-2* plants were comparable to those of WT plants (**Figure 2A,B**). *A_n_* values of *clpc2-2* leaves were lower than those of WT and *clpc2-2 promCLPC2:CLPC2* plants, but higher than those of *clpc1-1* plants (**Figure 2C**). Moreover, *g_s_* and *E* values of *clpc2-2* leaves were lower than those of WT plants (**Figure 2C**). Collectively, the data further indicated functional specificities between CLPC1 and CLPC2 and provided evidence that the reduced *A_n_* observed in *clpc1-1* in plants could be due to impairments in energy production during the light phase of photosynthesis, whereas the reduced *A_n_* in *clpc2-2* plants could be due, at least in part, to reduced CO_2_ diffusion through stomata.

**Figure 2:**
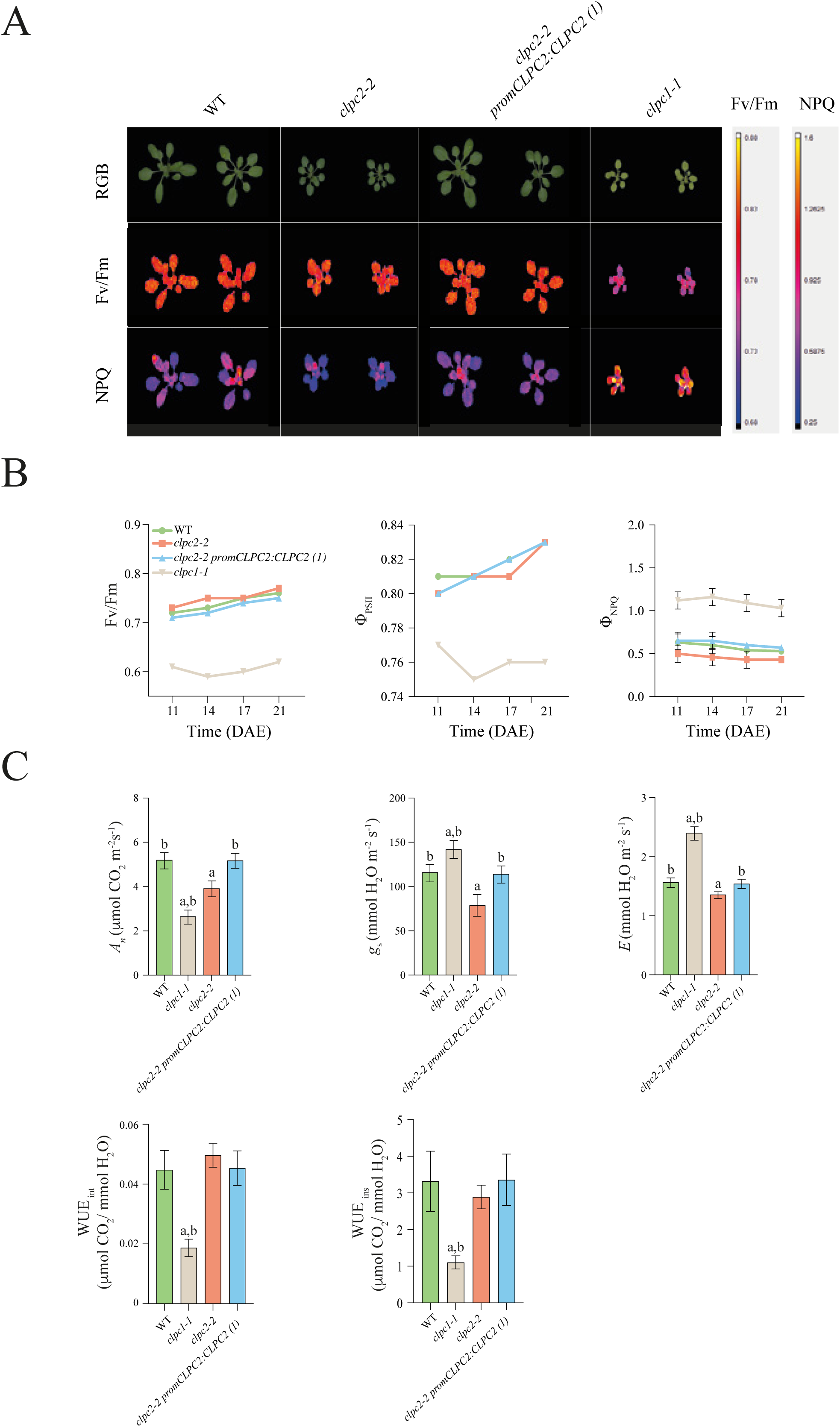
CLPC1 and CLPC2 mediate photosynthesis through different action mechanisms. (A) Fluorescence imaging of 21 days after sowing WT, *clpc2-2, clpc1-1* plants and plants of one representative line (line 1) of *clpc2-2 promCLPC2:CLPC2* grown on soil. (B) Time-course of *Fv/Fm*, <λ_PSII_ and <λ_NPQ_ in WT, *clpc2-2, clpc1-1* plants and plants of one representative line (line 1) of *clpc2-2 promCLPC2:CLPC2* grown on soil. (C) Graphics representing net CO_2_ uptake (*A_n_*), stomatal conductance (*g_s_*), transpiration (*E*) and intrinsic and instantaneous water use efficiencies (WUE-int and WUE-ins, respectively) in source leaves of 21 days after sowing WT, *clpc2-2* and *clpc1-1* plants, and plants of one representative line of *clpc2-2 promCLPC2:CLPC2* grown on soil. In “B”, values represent the mean ± SE of two independent experiments using 5 plants per genotype in each experiment. In “C”, values represent the means ± SE determined from three independent experiments using five plants per genotype in each experiment. Letters “a” and “b” indicate significant differences, according to Student’s t-test (P<0.05), between: “a” WT and other plants, and “b” *clpc2-2* and other plants. “DAE”, days after establishment.

### Knocking out *CLPC2* promotes hormonal changes that can account for the reduced photosynthesis and growth of *clpc2-2* plants

To investigate possible reasons for the reduced photosynthesis and slow growth phenotypes of *clpc2-2* plants, we conducted hormonomic analyses of WT and *clpc2-2* leaves. As shown in **Figure 3**, the lack of CLPC2 caused an increase in the levels of salicylic acid (SA), cis-12-oxophytodienic acid (cis-OPDA) and abscisic acid (ABA). These stress-related hormones play important roles in preventing the accumulation of excess reactive oxygen species and the consequent oxidative damages (Mateo et al., 2006; Bali et al., 2023). Furthermore, they strongly determine photosynthesis and growth as they negatively regulate stomatal opening and CO_2_ availability (Mateo et al., 2006; González-Guzmán et al., 2012; Savchenko et al., 2014). Therefore, the reduced *g_s_* and *E* values of *clpc2-2* leaves could be ascribed, at least in part, to high levels of ABA, SA and cis-OPDA. The lack of CLPC2 also caused a decrease in the levels of indole acetic acid (IAA) and the total content of transport and active forms of plastidic-type isopentenyladenine (iP) and *trans*-zeatin (tZ) and cytosolic-type *cis-*zeatin CK. CK positively regulates photosynthesis and growth (Cortleven and Valcke, 2012; Kieber and Schaller, 2014). It thus appears that (i) CLPC2 mediates growth and photosynthesis by regulating hormone balance and signalling and (ii) the reduced photosynthesis and growth of *clpc2-2* is due, at least partly, to high levels of the stress-related hormones cis-OPDA, ABA and SA and reduced levels of active CK.

**Figure 3:**
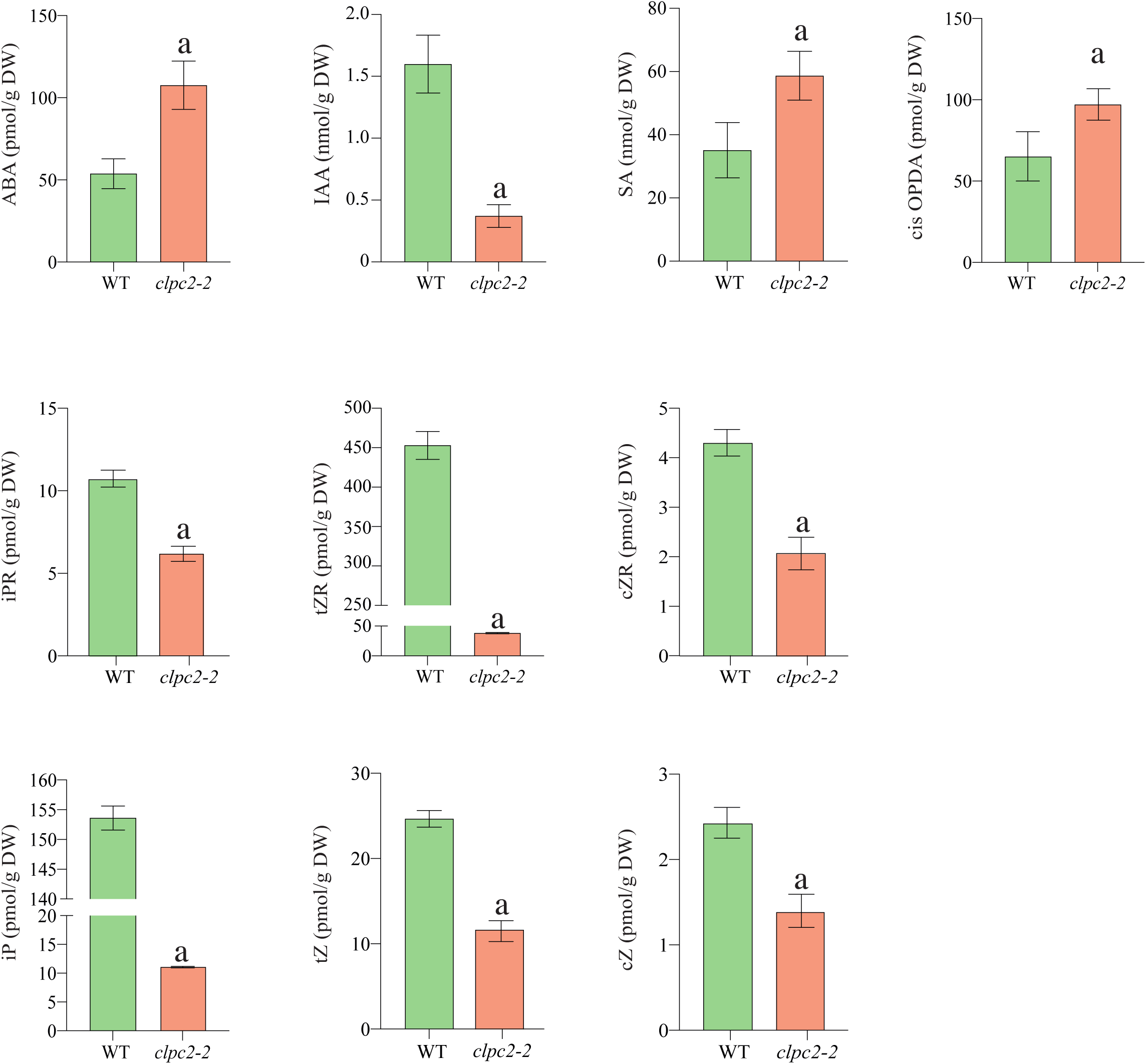
Knocking out *CLPC2* promotes changes in the leaf hormonome. Contents of the indicated hormones in leaves of 20 days after sowing WT and *clpc2-2* plants. Values represent the means ± SE determined from three replicates of 10 plants. Asterisks (*) indicate significant differences based on Student’s t-tests (P<0.05).

### CLPC2 is an important determinant of embryo development

Previous studies on the role of CLPC1 and CLPC2 in seed development showed that developing siliques of single *clpc1* and *clpc2* knockout mutans had seeds and ovules with WT external phenotype (Kovacheva et al., 2007), whereas siliques of self-pollinated heterozygous double mutants had seeds with abnormal phenotypes. Attempts to identify double *clpc1clpc2* knockout mutants were unsuccessful, strongly indicating that the double *clpc1clpc2* knockout genotype causes embryo lethality and that the two chaperones are partially redundant (Kovacheva et al., 2007). However, the phenotypes of mature seeds of single knockout *clpc1* and *clpc2* mutants were not characterized. Therefore, it was not possible to determine which one of the two chaperones has a dominant role in seed development. Here we found that dry *clpc2-2* seeds were 33% lighter than WT, *clpc2-2 promCLPC2:CLPC2* and *clpc1-1* seeds (**Figure 4A**). Furthermore, unlike WT, *clpc2-2 promCLPC2:CLPC2* and *clpc1-1* seeds, *clpc2-2* seeds displayed a wrinkled phenotype (**Figure 4B**). *clpc2-2* seeds had germination rates 20% lower than those of WT and *clpc1-1* seeds on MS agar with no supplemental sugar (**Figure 4C,D**) (**Supplemental Table 1**). Moreover, establishment rates of autotrophically grown *clpc2-2* seedlings were ca. 10% those of WT and *clpc1-1* seedlings (**Figure 4C,D**) (**Supplemental Table 1**). Both germination and establishment rates of *clpc2-2* substantially increased when sucrose was included in the culture medium (**Figure 4C,D**) (**Supplemental Table 1**). Once transferred to soil, heterotrophically grown *clpc2-2* seedlings developed, flowered and produced viable seeds. These findings showed that CLPC2 is an important determinant of seed growth, germination and seedling establishment.

**Figure 4.**
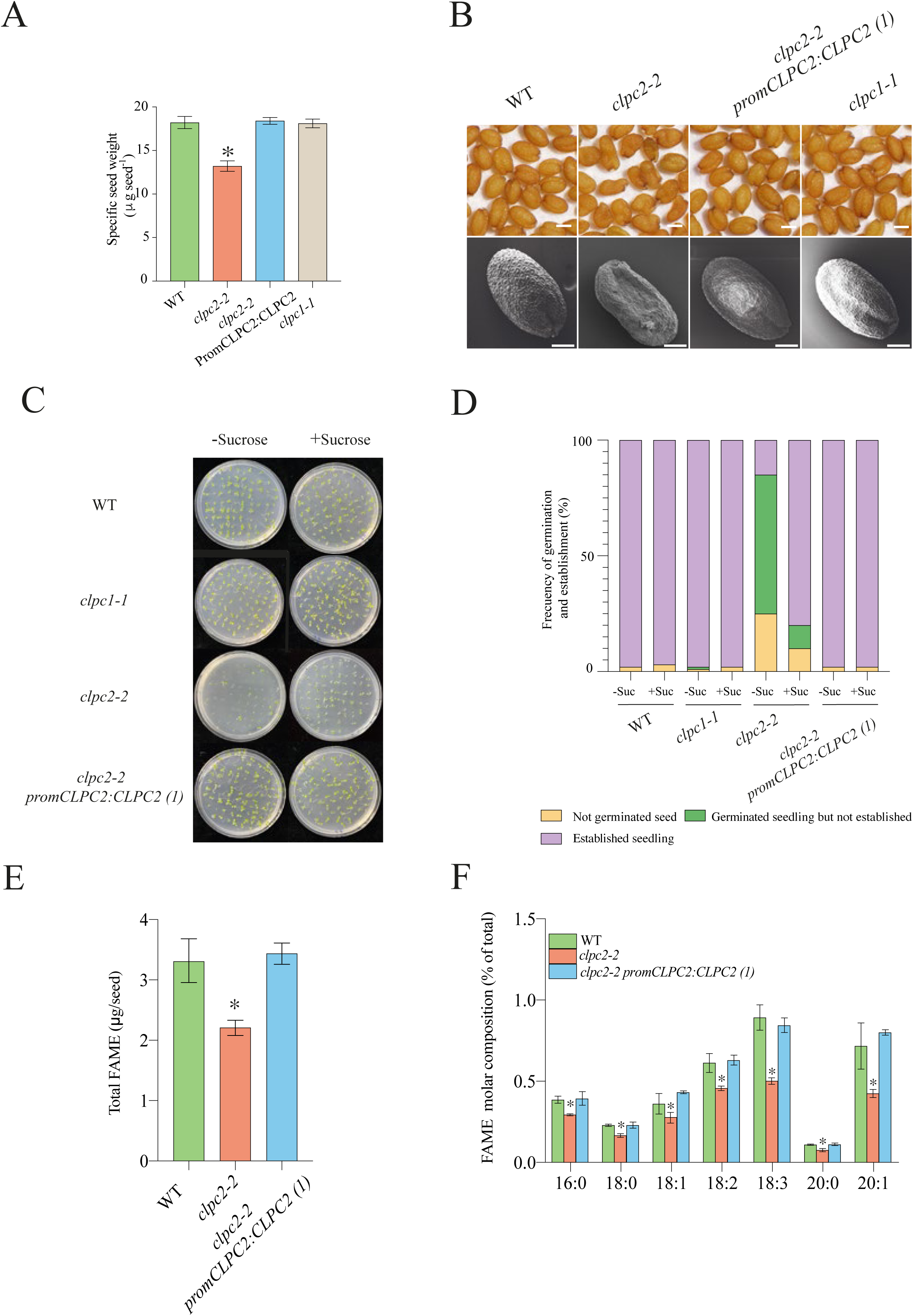
CLPC2 is an important determinant of seed yield and composition and seedling establishment. (A) Specific weight and (B) external phenotype (top panel) and scanning electron micrographs (bottom panel) of seeds from WT, *clpc2-2* and *clpc1-1* plants and one representative line (line 1) of *clpc2-2 promCLPC2:CLPC2* plants. (C) Photographs and (D) germination and establishment rates of WT, *clpc2-2, clpc1-1* and *clpc2-2 promCLPC2:CLPC2(1)* plants (8 days after sowing) grown on solid half-strength MS medium with or without sucrose (Suc) supplementation. (E) Total fatty acid methylester (FAME) contents per seed and (F) FAME profile of dry mature WT, *clpc2-2* and *clpc2-2 promCLPC2:CLPC2(1)* seeds. Values in (D) represent the means ± SE determined from a representative case of three biological replicates, each replicate being ca. 80 plants (data shown in **Supplemental Table 1)**. Values in (A), (E) and (F) represent the means ± SE of three biological replicates, each biological replicate being a pool of 20 seeds. Asterisks indicate significant differences from WT seeds according to Student’s t-tests (*p*<0.05). Scale bars in (B) = 200 μm (top panel) and 100 μm (bottom panel).

Oil is a major component of dry Arabidopsis seeds, which is used to support postgerminative seedling growth and establishment. Seeds of mutants with defects in FA synthesis display a wrinkled phenotype and have reduced establishment rates in the absence of an exogenous sugar source (Baud et al., 2008; Bahaji et al., 2018). This suggested that *clpc2-2* seeds could have reduced lipid content. In line with this presumption, analyses of FA levels (following methylesterification) showed that *clpc2-2* seeds accumulated 35% less oil than WT and *clpc2-2 promCLPC2:CLPC2* seeds at maturity (**Figure 4E**). Subsequent FA composition analyses revealed that, compared with WT and *promCLPC2:CLPC2* seeds, *clpc2-2* seeds have reduced relative contents of all FA forms including saturated FAs such as palmitic acid (16:0), stearic acid (18:0) and arachidic acid (20:0), their desaturation intermediates, oleic (18:1) and linoleic (18:2) acids, and their end products of desaturation and elongation, linolenic (18:3) and eicosanoic (20:1) acids (**Figure 4F**). This relative FA content profile contrasts with those of low seed oil mutants impared in FA biosynthesis, characterized by having relatively higher than WT levels of linolenic and eicosanoic acids and WT levels of palmitic and stearic acids (Baud et al., 2007; Bahaji et al., 2018). Therefore, the wrinkled phenotype of *clpc2-2* seeds is not directly due to reduced activity of FA biosynthesis pathways.

Some mutants of the CLPPR core produce wrinkled seeds and have reduced germination and establishment rates when grown under autotrophic conditions due to aberrant embryo growth and delayed development (Kim et al., 2009; Kim et al., 2013). This developmental arrest can be rescued by adding sucrose to the culture medium. We addressed the possibility that, like in *clppr* mutants, the wrinkled seed phenotype and reduced germination and establishment rates of *clpc2-2* plants could be related to impairments in embryo development.

As shown in **Figure 5A**, developing *clpc2-2* siliques had aberrant and aborted seeds at 15 days after pollination (DAP). Analyses of embryonic phenotypes in a series of developing pollinated pistils of *clpc2-2* plants revealed delayed embryogenesis (**Figure 5B** and **C**) and reduced number of nuclei in endosperm cells (**Figure 5D** and **E**), strongly indicating that the wrinkled phenotype of *clpc2-2* seeds is due to aberrant embryo growth and delayed development. This developmental defect was not fully penetrant across all seed sets. In *clpc2-2* plants, aberrant phenotypes were observed in approximately 20% of seeds within developing siliques at 15 DAP, including a residual number of ovules and seed abortions (**Supplemental Figure 2A**). At maturity, *clpc2-2* seeds displayed a major impact in phenotypic severity, categorized as “strongly aberrant” (∼5%), “aberrant” (∼35%), and “mild aberrant” (∼40%), affecting around 80% of seeds per silique (**Supplemental Figure 2B,C**). Collectively, our data show that (i) CLPC2 is an important determinant of embryo development likely due to its involvement in the CLP complex-mediated plastidial protein homeostasis, acting as a major driver of seed expansion from the earliest stages by synchronizing control of the morphogenetic programs of both the embryo and the endosperm and (ii) CLPC2 is dominant over CLPC1 in this trait.

**Figure 5:**
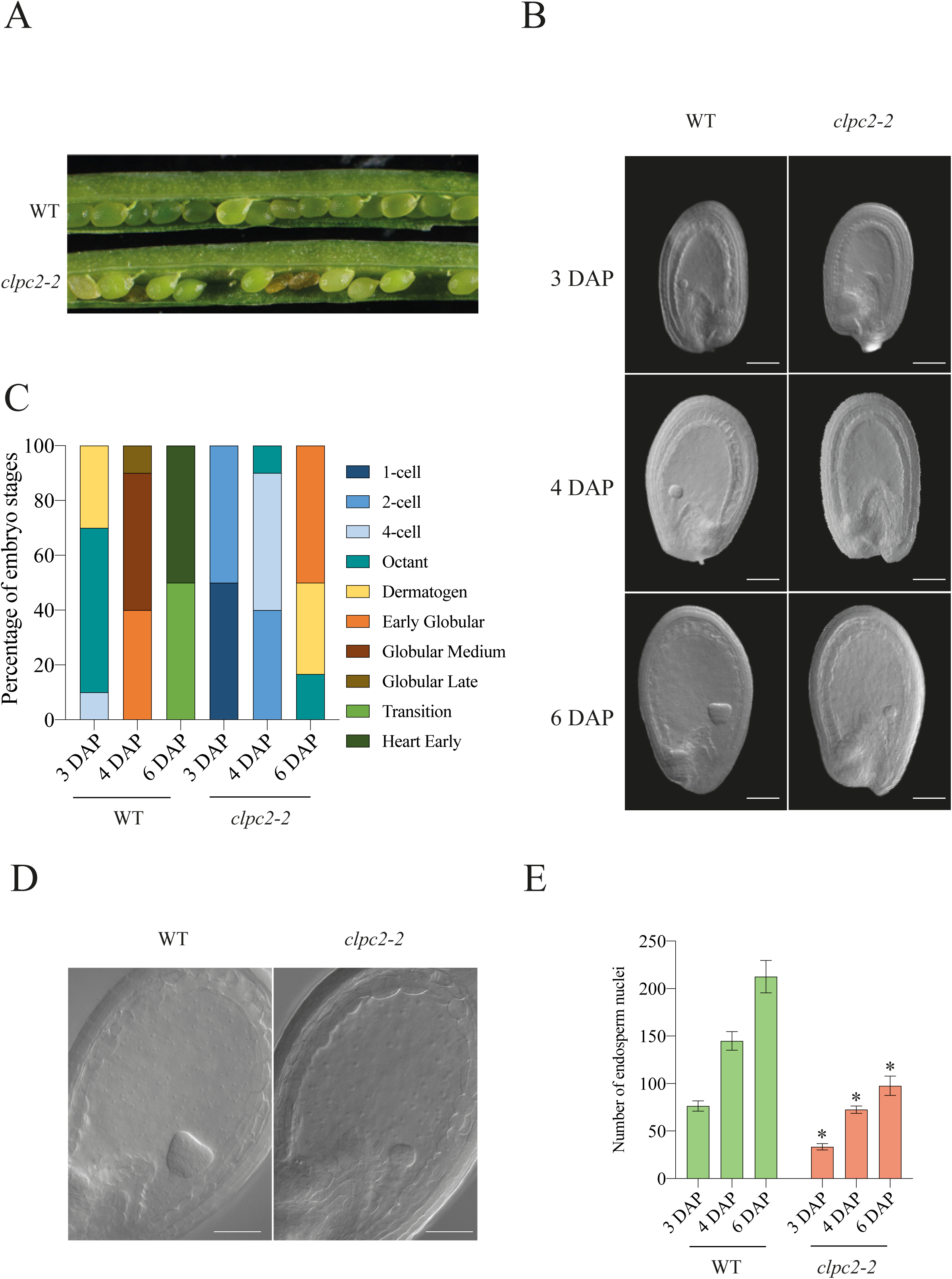
CLPC2 is an important determinant of embryo development. (A) Representative siliques of WT and *clpc2-2* plants at 15 days after pollination (DAP). (B) Scaning electron microscopy and (C) quantification of developmental stages of WT and *clpc2-2* seeds at 3, 4, and 6 DAP. Note that at 3 DAP, *clpc2-2* seeds had more embryos in preliminary developmental stages (1- and 2- cell) than WT seeds, which were mostly at the dermatogen and octant stages. At 4 DAP, *clpc2-2* seeds were at the 2- and 4-cell stages, while WT seeds had progressed to the globular medium and early stages. At 6 DAP, *clpc2-2* seeds were mostly at the early globular and dermatogen stage, whereas WT seeds had reached the transition and heart stages. (D) Scaning electron microscopy analysis for endosperm cellularization and (E) quantification of nuclei progression in WT and *clpc2-2* seeds at 3, 4, and 6 DAP. Note that in WT seeds the endosperm undergoes rapid nuclear divisions around 3 DAP, whereas in *clpc2-2* seeds endosperm development is delayed, with reduced nuclear proliferation. Scale bars: 50 μm (B), 50 μm (D). Values in (E) represent the means ± SE of 10 seeds per genotype, sampled from three independent siliques. Statistical significance was assassed using one-way ANOVA followed by Tukey’s HSD text.

### The CLPC2 interactome is distinct from that of CLPC1 but partially resembles to that of CLPPR

The above results showing differences in growth responses to fungal VCs, photosynthesis, seed development and seedling establishment between *clpc1-1* and *clpc2-2* plants, highlighted contrasting functional specificities for CLPC1 and CLPC2. This would indicate that CLPC1 and CLPC2 differ in their molecular mechanisms of action, including substrate recognition specificity. To test this hypothesis, we first conducted high-throughput differential proteomic analyses between *clpc2-2* and WT leaves. We found that 79 and 34 of the 5451 proteins identified in this study were significantly up- and down-regulated, respectively, by the lack of CLPC2 (**Table 1**, **Supplemental Table 2**). As expected, CLPC2 was identified in WT leaves, but not in *clpc2-2* leaves. Twenty-seven of the 113 differentially expressed proteins (DEPs) had a predicted plastidial localization, and 20 of them were up-regulated in *clpc2-2* leaves (**Table 1**). RT-qPCR analyses of the levels of transcripts encoding some of these plastidial proteins showed that *clpc2-2* leaves accumulate WT or lower than WT levels of these transcripts (**Supplemental Figure 3**). Therefore, we hypothesize that the increased levels of some of these plastidial proteins in *clpc2-2* leaves could be due, at least in part, to a longer lifespan caused by lower CLP protease activity.

**Table 1:**
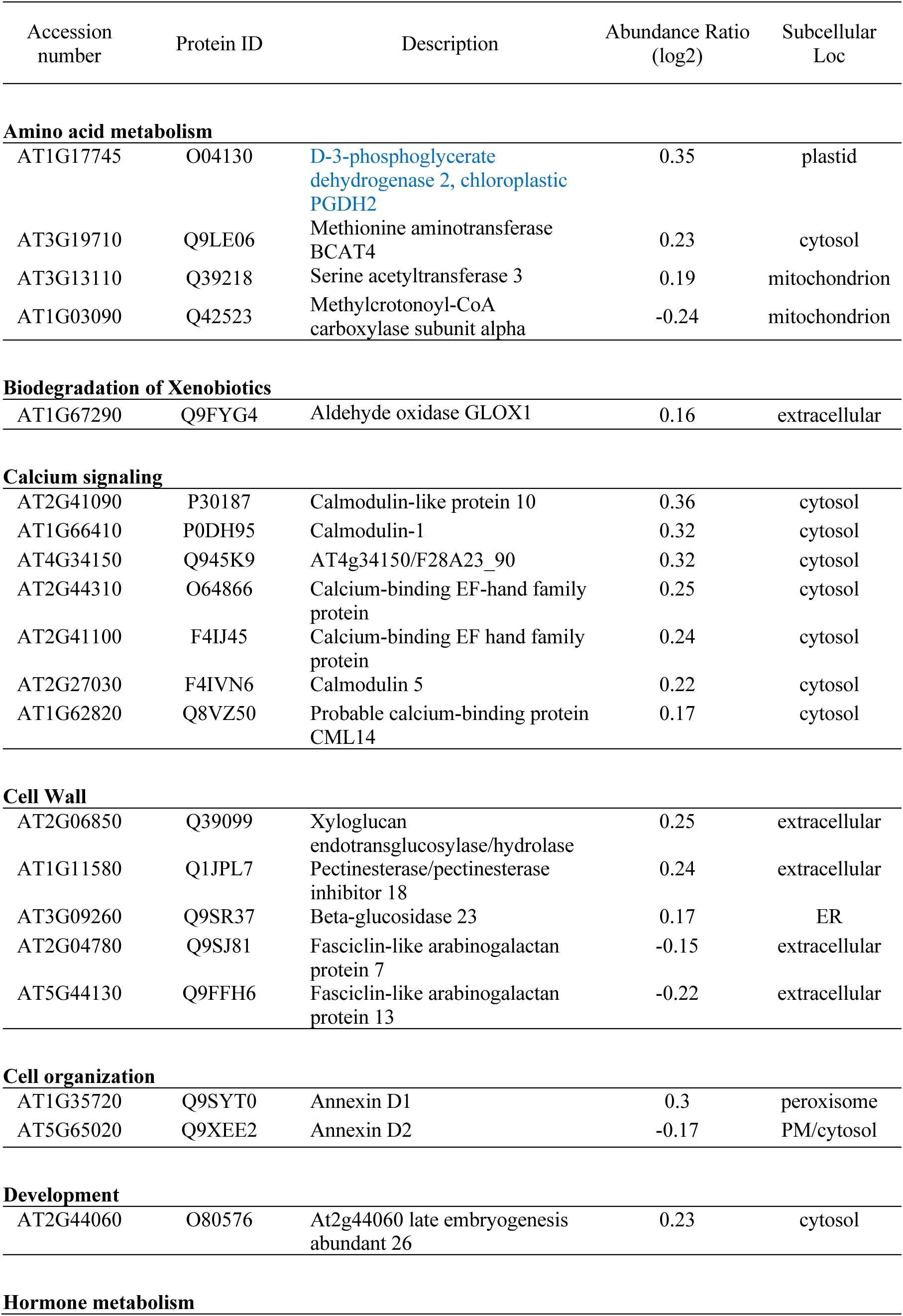
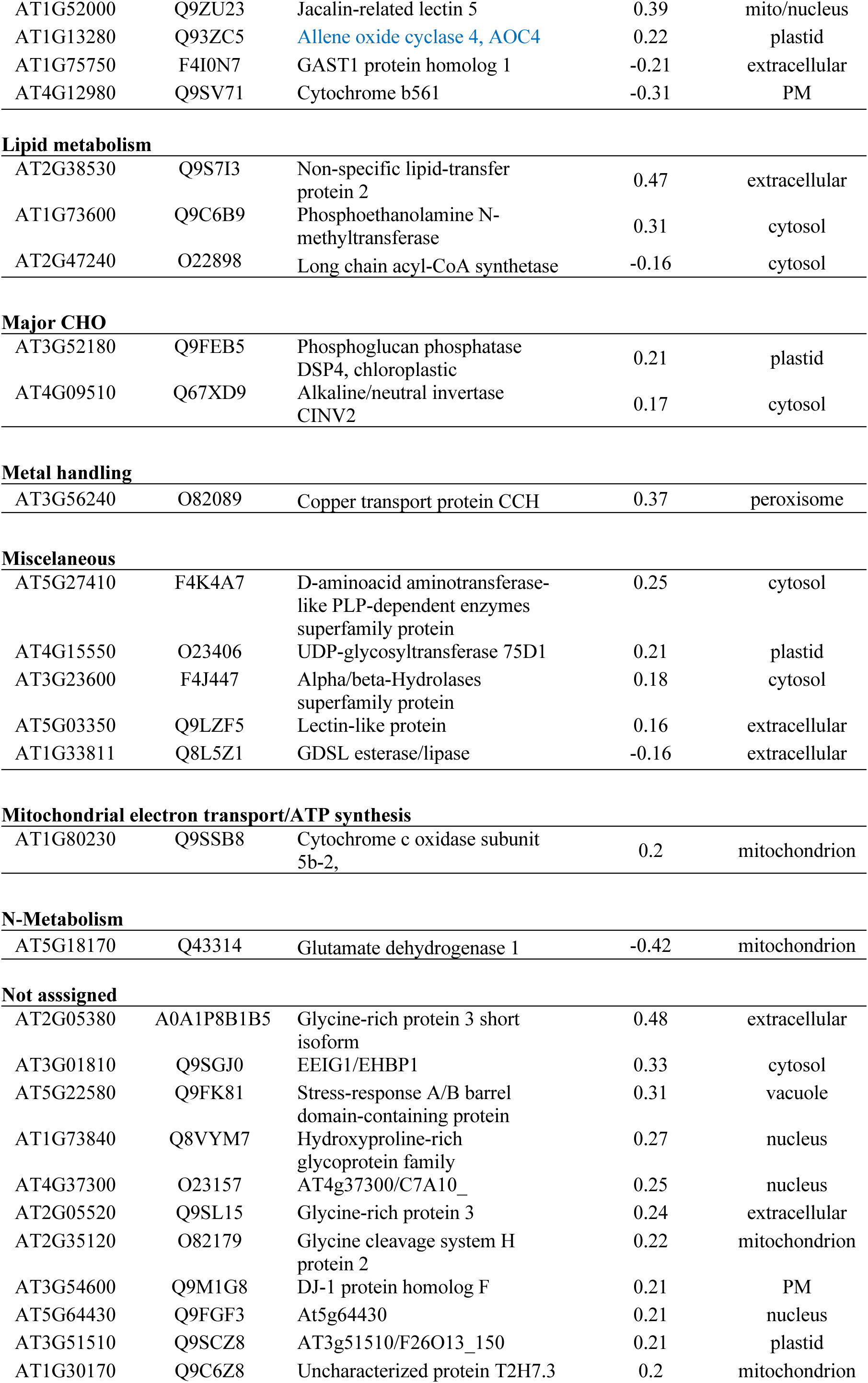
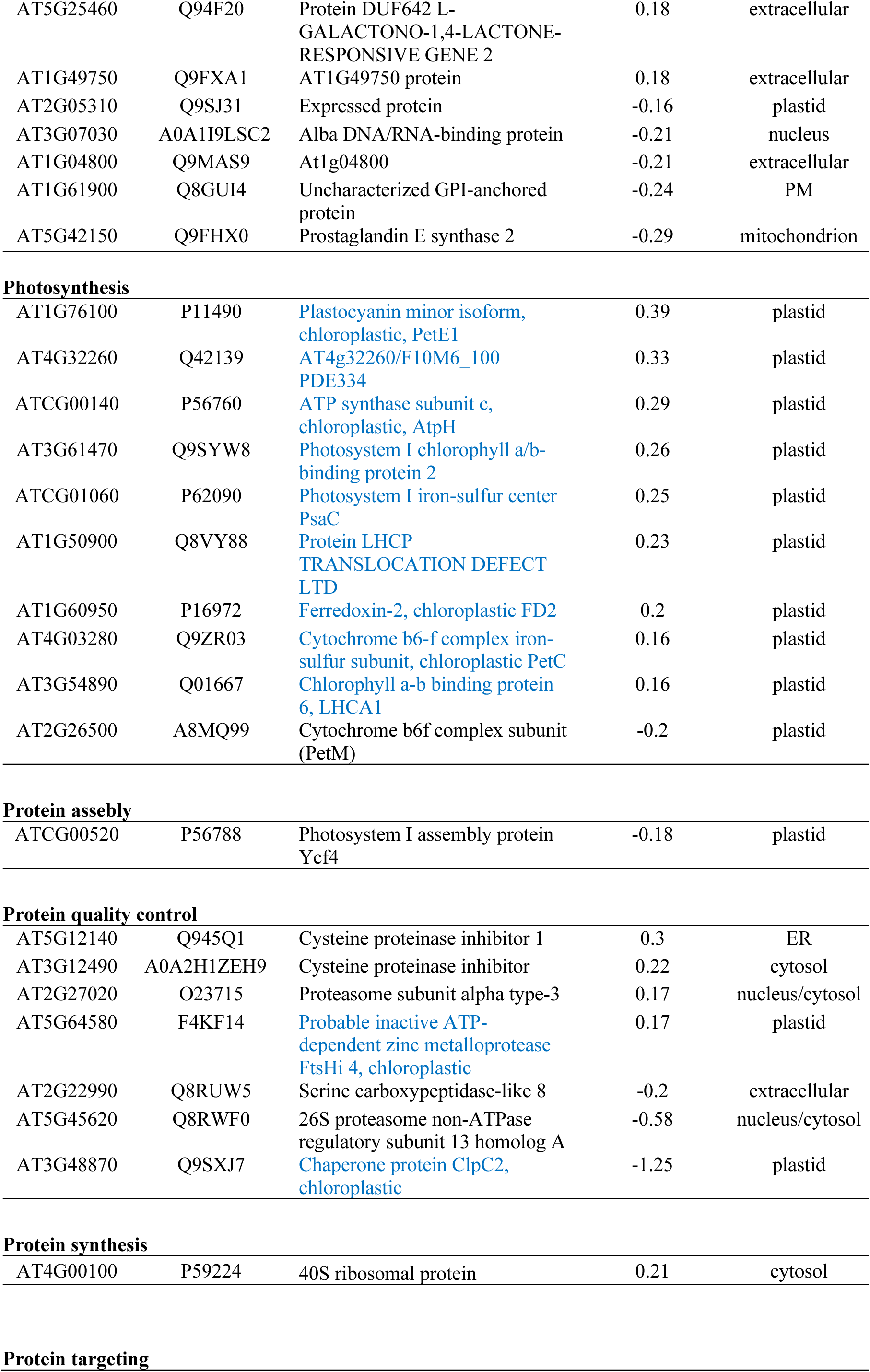
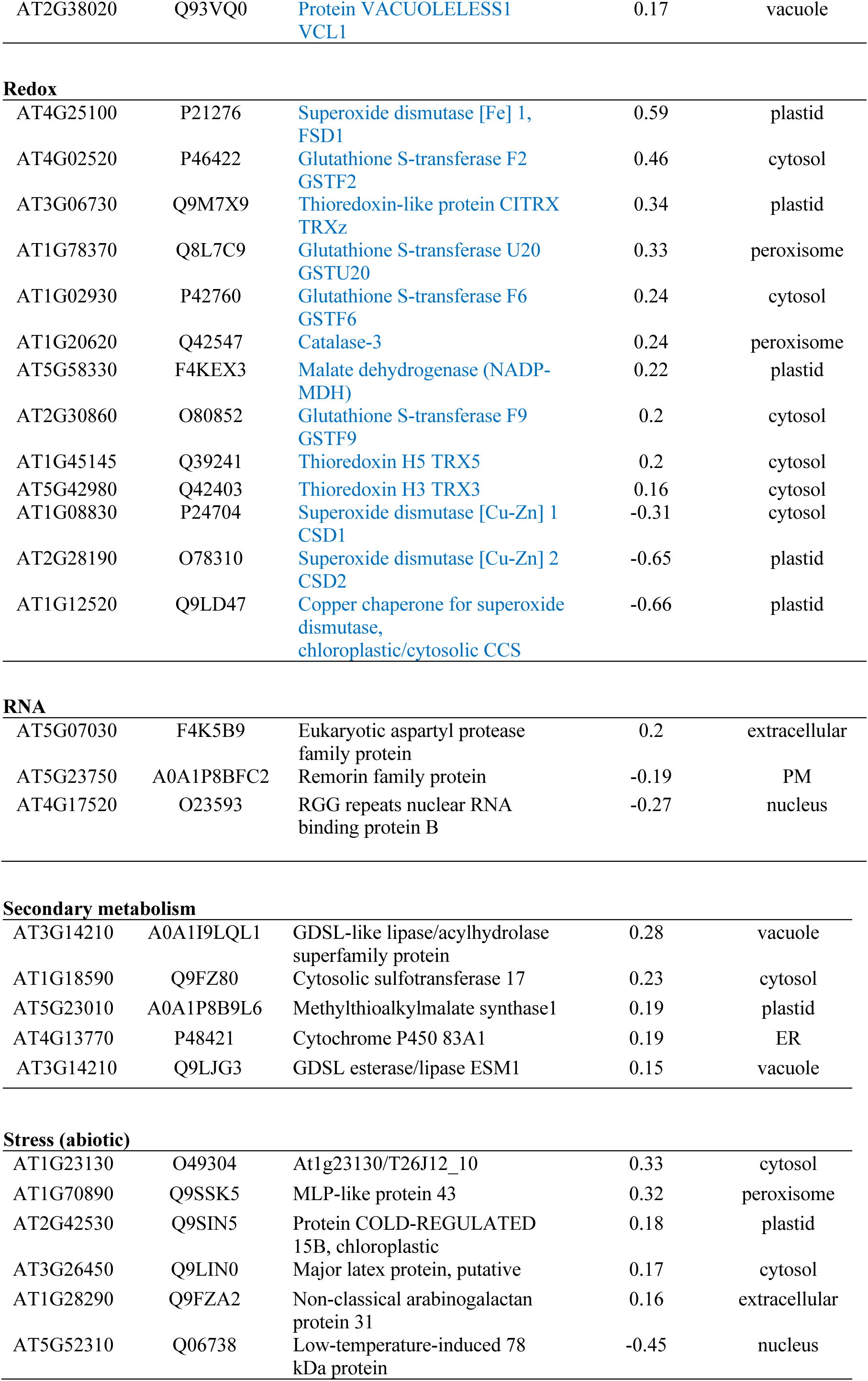
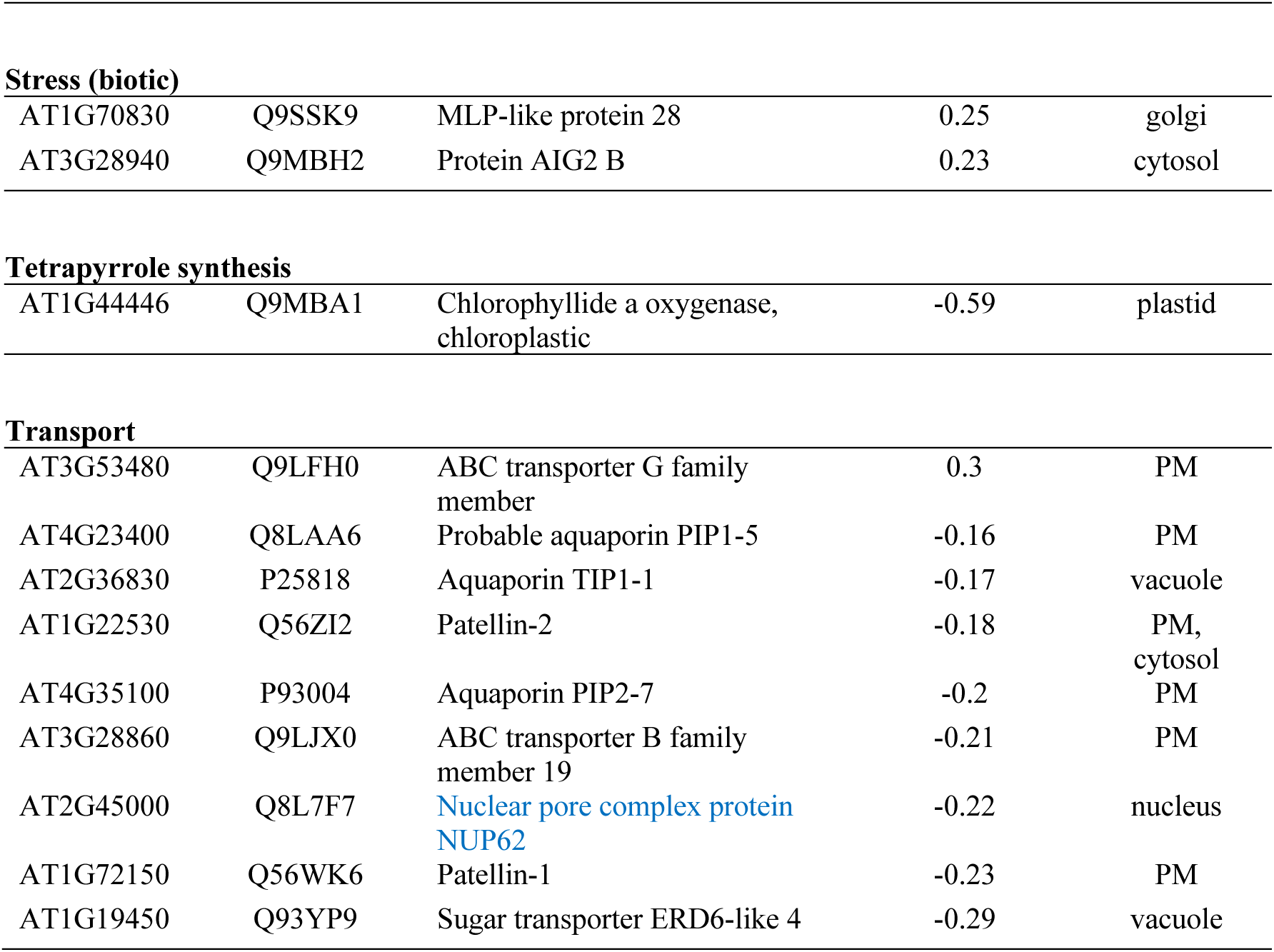
List of proteins differentially accumulated by the lack of CLPC2 in leaves (*clpc2-2* vs. WT). Proteins are classified according to their functions. Proteins discussed in the main text are highlighted in blue.

Studies on the biological processes affected by the lack of CLPC2 using the MapMan tool (https://mapman.gabipd.org/) revealed that the 113 proteins differentially accumulated in *clpc2-2* leaves are involved in multiple processes (**Figure 6**, **Table 1**). Some of these proteins play important roles in amino acid metabolism (e.g. PGDH2), hormone metabolism (e.g. AOC4), nitrogen metabolim (e.g. GDH1), photosynthesis (e.g. PetE1, PDE334, AtpH, LHCA1, LHCA2, PsaC, LTD, FD2 and PetC), protein quality control (e.g. FtsHi4) and targeting (e.g. VCL1), redox regulation and oxidative stress (e.g. FSD1, GSTF2, TRXz, GSTU20, GSTF6, CAT3, GSTF9, TRX5, TRX3, CSD1, CSD2 and CCS), energy homeostasis (e.g. NADP-MDH) and transport (e.g. NUP62) (**Figure 6**, **Table 1**). From these, the following plastidial proteins overaccumulating in *clpc2-2* leaves play important roles in embryo development, seedling establishment and growth: TRXz, which is a thioredoxin that is essential for germination and seedling establishment and growth (Meng et al., 2010; Arsova et al., 2010); FtsHi4, which is a pseudoprotease that forms part of a FtsH12-FtsHi complex involved in ATP-driven protein import across the chloroplast envelope that is essential for chloroplast development and embryogenesis (Lu et al., 2014; Kikuchi et al., 2018; Schreier et al., 2018; Meinke, 2019); petC, which is a subunit of the cytochrome b6-f complex that is essential for photosynthesis and growth (Maiwald et al., 2003); FD2, which is the major form of ferredoxins that together with antioxidant defences such as NADP-MDH and FSD1, plays an important role in plastidial NADPH homeostasis and growth, and in plant protection against oxidative damage in response to environmental changes (Liu et al., 2013; Dvorák et al., 2020; Yokochi et al., 2021). We hypothesize that the reduced establishment and growth, as well as the delayed embryonic development of *clpc2-2* plants, may be attributed, at least in part, to impaired CLPC2-mediated processing of some of these plastidial proteins. Altered levels of extraplastidial proteins essential for growth and embryogenesis, such as NUP62 (Parry, 2014; Boeglin et al., 2016; Meinke, 2019) and VCL1 (Rojo et al., 2001; Meinke, 2019) could also explain the phenotype of *clpc2-2* plants.

**Figure 6:**
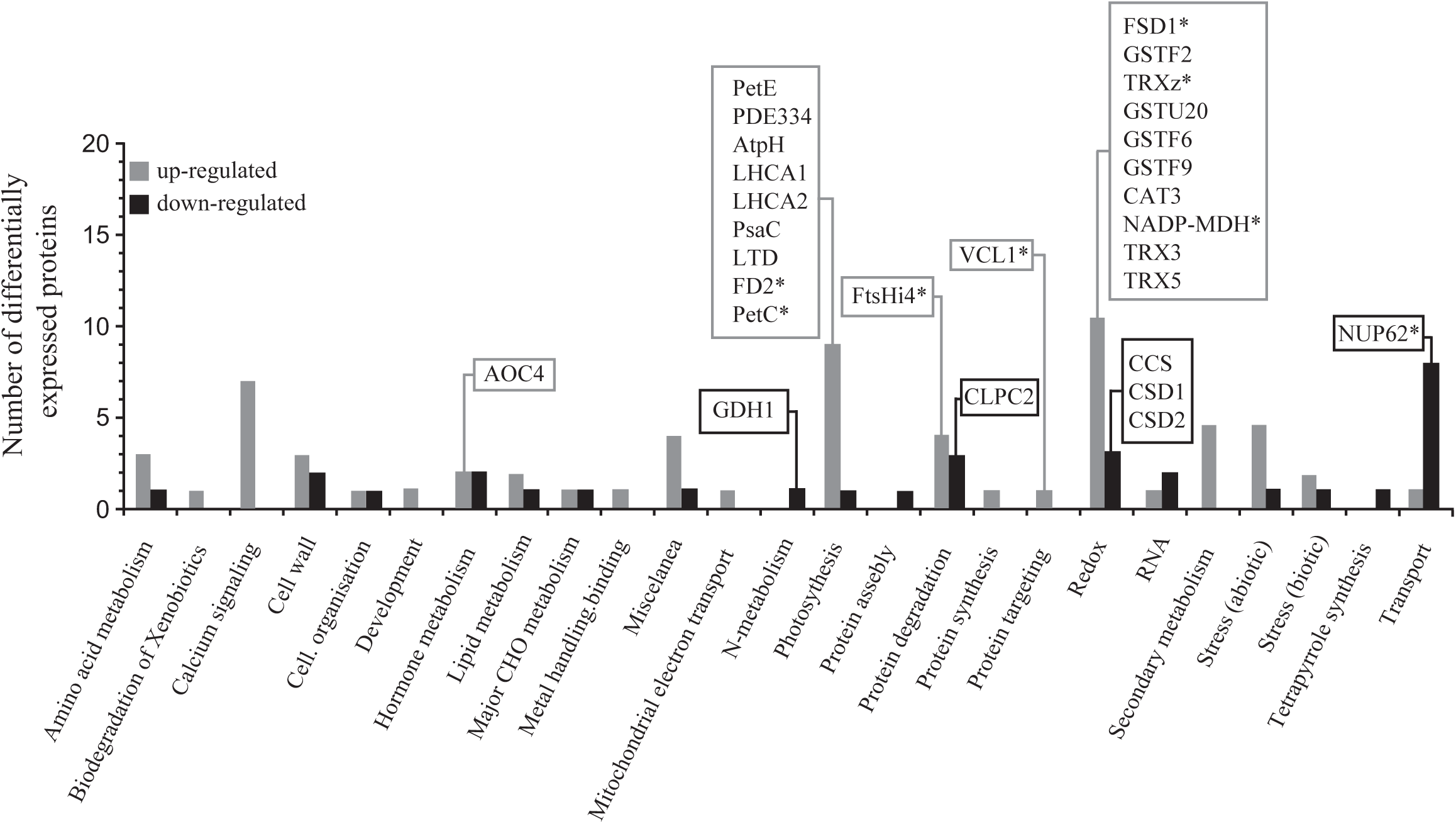
Knocking out *CLPC2* promotes changes in the leaf proteome. The graphic represents the functional categorization according to the Mapman tool of the proteins that are differentially accumulated in 20 days after sowing *clpc2-2* and WT leaves (*clpc2-2* vs. WT). The numbers of up- and down-regulated proteins in each categorical group are indicated by gray and black bars, respectively. The data were obtained from **Table 1**. Proteins discussed here are boxed and proteins that play important roles in embryo development, seedling establishment and growth are indicated with asterisks.

To obtain further insight into the molecular action mechanisms of CLPC2 and possible similarities with those mediated by CLPC1 and CLPPR, we compared the proteomic phenotype of *clpc2-2* leaves with those of *clpr2-1, clpr4-1* and *clpp3-1* proteolytic CLPPR core mutants (Kim et al., 2009; Zybailov et al., 2009; Kim et al., 2013) (**Supplemental Table 3**) and *clpc1-1* chloroplasts (cf. Supplemental Table 3 in Nishimura et al., 2013). We also compared the list of 59 CLPC1 interactors identified by Liao et al. (2022) with the list of 113 proteins differentially accumulated by the lack of CLPC2. Except for CLPC2, none of the proteins differentially accumulated by the lack of CLPC2 matched any of the CLPC1 interactors identified by Liao et al. (2022). Furthermore, only 2 of the 108 chloroplastic proteins differentially accumulated by the lack of CLPC1 were also differentially accumulated by the lack of CLPC2 (e.g. PetC and CCS), strongly indicating that CLPC1 and CLPC2 interactors are different. Notably, fifteen of the 309 leaf proteins differentially accumulated by the lack of CLPPR subunits (e.g. GSTF2, GSTF9, TRXz, PDE334, MLP43, ANN1, ESM1, BCAT4, At3g26450, petC, LHCA1, LLP, GDH1, RD29A, CCS and CLPC2) were also differentially accumulated by the lack of CLPC2 (**Supplemental Table 3**). This would indicate that CLPC2 operates, at least in part, through CLPPR action (see below).

Next, we compared the proteomic phenotypes of *clpc1-1, clpc2-2* and *clppr* leaves with those of WT plants exposed to fungal VCs (cf. Supplemental Table 3 in Ameztoy et al., 2021). We found that many proteins differentially accumulated by the lack of CLPC1, CLPC2 and CLPPR were VC-responsive, strongly indicating that the proteomic response of plants to fungal VCs involves CLP complex action. Thus, as shown in **Supplemental Table 4** and **Supplemental Table 5**, 44 of the 108 plastidial proteins differentially accumulated by the lack of CLPC1 and 43 of the 113 proteins differentially accumulated by the lack of CLPC2 were VC-responsive, respectively. Furthermore, 125 of the 309 proteins differentially accumulated by CLPPR inactivation were also differentially accumulated by the VC treatment, including CLPC2 (**Supplemental Table 6**). Whereas VC-responsive proteins that are differentially expressed by the lack of CLPC1 mainly comprise MEP pathway and chlorophyll biosynthesis enzymes and proteins involved in chloroplast proteostasis (**Supplemental Table 4**), VC-responsive proteins that are differentially expressed by the lack of CLPC2 are involved in other processes (**Supplemental Table 5**), strongly indicating that the response of plants to fungal VCs is largely mediated by distinct CLPC1- and CLPC2-specific mechanisms. Collectively, the data would strongly indicate that CLPC1 and CLPC2 interactors are different and both chaperones regulate different processes, at least partly, through CLP-mediated action.

### CLPC2 overexpression causes variegation and promotes changes in the proteome similar to those promoted by the lack of CLPC1

CLPC1 and CLPC2 function both as hexameric regulatory subunits of the CLP protease complex and independently as dimeric chaperones (Peltier et al., 2004). CLPC2 can be recognized by the *in vivo* CLPC1 substrate trapping CLPC1-TRAP system (Montandon et al., 2019). This and the facts that (i) leaves of CLPPR core mutants accumulate higher than WT levels of CLPC2 (Kim et al., 2009; Zybailov et al., 2009; Kim et al. 2013), (ii) the proteomic profiles of *clpc2-2* and *clppr* leaves partially overlap (this work), and (iii) some phenotypic characteristics of *clpc2-2* and *clppr* plants are similar (this work) would indicate that (i) CLPC1 and CLPC2 compete with each other within the hexameric regulatory chaperone complex to bind to the proteolytic CLPPR core and (ii) CLPC2 works, at least partly, through the CLP complex. To test these hypotheses, we carried out immunoblot analyses of CPN20, ZEP and DXS in leaves of on soil grown WT, *clpc1-1* and *clpc2-2* plants, and WT plants expressing *CLPC2* under the control of the 35S promoter (35S:CLPC2 plants). These plastidial proteins are clients of CLPC1 and are known to overaccumulate in *clpc1-1* plants (Nishimura et al., 2013; Liao et al., 2022). We reasoned that if CLPC1 and CLPC2 have different substrate specificites, and both chaperones compete for the CLPPR core, levels of CPN20, ZEP and DXS in 35S:CLPC2 plants should be higher than in WT plants. Conversely, if CLPC2 and CLPC1 do not compete to form part of the CLP complex, 35S:CLPC2 leaves should accumulate WT levels of these proteins.

In accordance with Kovacheva et al. (2007), the 35S:CLPC2 plants showed a dwarf phenotype and had chlorotic, pale-green leaves which sporadically presented white sectors (**Figure 7A**). FtsH mutants impaired in thylakoid membrane biogenesis and maintenance show leaves with white sectors (Chen et al., 2000; Park and Rodermel, 2004; Zaltsman et al., 2005). This variegated phenotype of FtsH mutants can be suppressed by mutations in *CLPC2* (Park and Rodermel, 2004) and in *CLPR1* (Yu et al., 2008). Therefore, it is likely that the white sectors of leaves of 35S:CLPC2 plants is due to FtsH malfunction caused by high CLPC2 activity. However, we cannot rule out the possibility that this phenomenon is due, at least in part, to silencing effects (Bazin et al., 2023). As expected, *clpc1-1* leaves accumulated higher than WT levels of CPN20, ZEP and DXS, whereas *clpc2-2* leaves accumulated WT-like levels of these proteins (**Figure 7B,C**). Notably, CPN20, ZEP and DXS levels in 35S:CLPC2 leaves were substantially higher than in WT leaves (**Figure 7B,C**). These findings strongly indicate that CLPC1 and CLPC2 can coexist in the same CLP complex, and further provide evidence that the two chaperones have different specificities toward chloroplast substrates.

**Figure 7:**
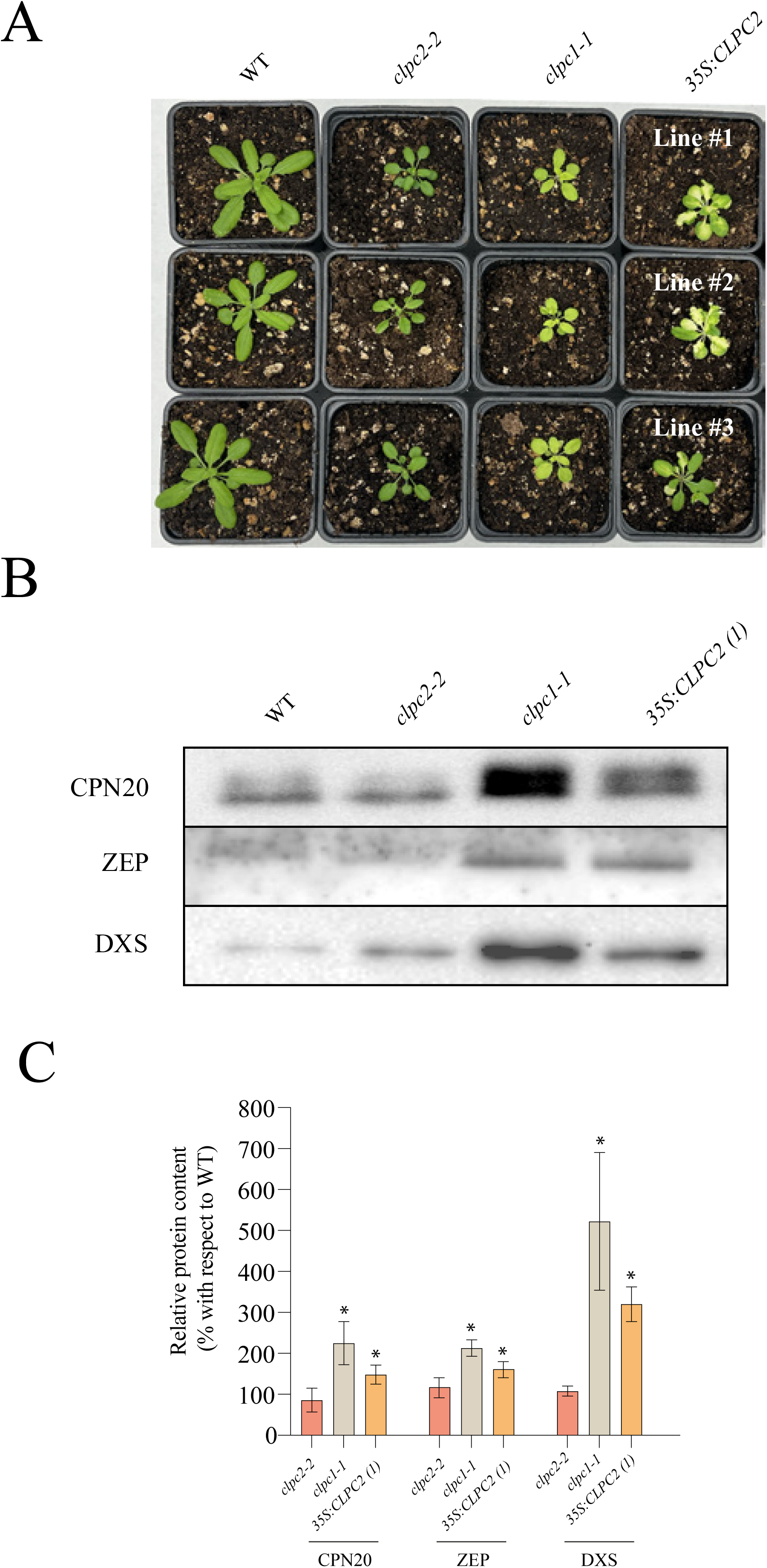
*35S* promoter-driven *CLPC2* expression promotes changes in the proteome similar to those promoted by the lack of CLPC1. (A) External phenotype of on soil-grown WT, *clpc1-1, clpc2-2* and three independent lines of WT plants expressing *CLPC2* under the control of the 35S promoter (35S:CLPC2 plants). (B) Immunoblots of protein samples extracted from leaves of WT, *clpc1-1, clpc2-2* plants and leaves of one representative line of 35S:CLPC2 plants. The blots were probed with antibodies against CPN20, ZEP and DXS. A representative example is shown. (C) CPN20, ZEP and DXS levels in leaves of *clpc1-1, clpc2-2* plants and leaves of one representative line of 35S:CLPC2 plants relative to those of leaves of WT plants. The means values ± SE from three independent experiments are shown. Asterisks above bars indicate significant differences according to Student’s t-test (P<0.05), between the WT and mutant plants. In (B), the gel was loaded with 60 μg of protein per lane.

## MATERIALS AND METHODS

### Plants, growth conditions and sampling

The work was carried out using *Arabidopsis thaliana* L. (Heynh) WT plants (ecotype Columbia Col-0), the *clpc2-2* (GABIKAT 039E12) and *clpc1-1* (SALK_014058) knockout mutants (López et al., 2024) obtained from the Nottingham Arabidopsis Stock Centre. In addition, we used three independent lines each of *clpc2-2* plants transformed with *promCLPC2:CLPC2*, which express CLPC2 under the control of 2.0 kbp immediately upstream of the translation start codon of CLPC2 (**Supplemental Figure 4, Supplemental Figure 5**), and WT plants expressing CLPC2 under the control of the cauliflower mosaic virus 35S promoter (*35S:CLPC2*) (**Supplemental Figure 4, Supplemental Figure 5**). To construct *promCLPC2:CLPC2* and *35S:CLPC2*, a complete cDNA corresponding to the CLPC2 gene (At3g48870) was obtained from the RIKEN Arabidopsis cDNA collection (pdy16624) (Seki et al., 1998; 2002), amplified by PCR using the specific ‘attB1 CLPC2’ and ‘attB2 CLPC2’ primers (**Supplemental Table 7**), cloned into the pDONR/Zeo plasmid using BP clonase (Invitrogen) (**Supplemental Figure 4**), and confirmed by sequencing. The plasmid constructs were transferred to *Agrobacterium tumefaciens* to transform plants according to Clough and Bent (1998). Plants were grown on soil or in Petri dishes (92 x 16 mm, Sarstedt, Ref. 82.1472.001) containing half-strength agar solidified Murashige and Skoog (MS) (Phytotechlab M519) medium with or without sucrose supplementation in growth chambers with a 16 h light (90 μmol photons sec^−1^ m^-2^ (22°C)/8 h dark (18°C) cycles. Fully expanded source leaves of plants grown on soil were harvested, and immediately frozen and ground in liquid nitrogen. To investigate effects of small fungal VCs emitted by the fungus *A. alternata* on plants we used the “plasticized PVC wrap and charcoal filter-based box-in-box” co-cultivation system (Gámez-Arcas et al., 2022a). *A. alternata* was grown in Petri dishes containing agar-solidified MS medium supplemented with 90 mM sucrose.

### Determination of photosynthetic parameters

Stomatal conductance (*g_s_*) and transpiration (*E*) values were determined using a LI-COR 6400 gas exchange portable photosynthesis system (LI-COR, Lincoln, NE, USA). Net rates of CO_2_ assimilation (*A_n_*) were calculated using equations developed by von Caemmerer & Farquhar (1981). Intrinsic and instantaneous water use efficiencies (WUE-int and WUE-ins, respectively) were calculated as the ratios of *A_n_* to *g_s_*, and *A_n_* to *E*, respectively, as described by Flexas et al. (2016). Chlorophyll fluorescence emission parameters were determined using a PlantScreen^TM^ Compact System (Photon Systems Instruments, Brno, Czech Republic) high-throughput phenotyping as described in Sánchez-López et al. (2016).

### Morphological analysis of seeds

Seeds were observed and photographed using an Olympus MVX10 stereomicroscope (Japan). Scanning electron microscopy analyses of gold-coated dry seeds were performed using a JEOL JSM-5610LV scanning electron microscope. To investigate seed size dynamics, embryo developmental stages, and endosperm formation between 3 and 6 days after pollination (DAP), seed clearing was performed following the protocol described by Loperfido et al. (2025). Carpels were fixed overnight in a 9:1 (v/v) ethanol:acetic acid solution and subsequently rehydrated through a graded ethanol series. Seeds were manually dissected from Arabidopsis siliques and incubated in a chloral hydrate:glycerol:water solution (8:1:3, w/v/v). Samples were immediately imaged using a Zeiss Axiophot D1 microscope equipped with differential interference contrast (DIC) optics and an Axiocam MRc5 camera (Zeiss), operated via Axiovision software (version 4.1). Endosperm nuclei were counted, and embryo developmental stages were classified and documented. Mature seed size was quantified using SmartGrain software.

### Analytical procedures

Protein content was determined using the Bradford method using Bio-Rad prepared reagent (Bio-Rad Laboratories, Hercules, CA, USA). Seed FA content and composition analyses were conducted as described in Bahaji et al. (2018). The endogenous levels of the different plant hormones, including ABA, IAA, SA, cis-OPDA, and several CKs, were measured according to the protocol described by Vrobel et al. (2024).

### Proteomic analysis

High-throughput, isobaric labeling-based differential proteomic analyses of 20 days after sowing WT and *clpc2-2* plants were conducted essentially as described in Gámez-Arcas et al. (2022b) using three biological replicates of 30 plants per replicate. Statistical significance was measured using *q*-values (FDR). The cut-off for identifying DEPs was established at FDR ≤ 0.05% and log2 ratios of > 0.15 (for proteins whose content was increased by the lack of CLPC2) or <-0.15 (for proteins whose content was reduced by the lack of CLPC2). Predicted subcellular localization studies of DEPs were conducted using the SUBA4 database.

### Western blot analyses

For immunoblot analyses, protein samples were separated on 10% SDS-PAGE, transferred to PVDF, and immunodecorated by using the antisera raised against CPN20, ZEP and DXS from PhytoAB (PHY2525S, PHY7274S, PHY0848S, respectively), and an HRP-conjugated antirabbit IgG (Agrisera) as a secondary antibody. The detection of the immunoreactive bands was performed using the ECL Select (Amersham). Chemiluminiscent signals were visualized using a ChemiDoc Imaging System (Bio-Rad) and quantified with ImageJ. Immunoblots from four independent experiments were quantified.

### Real-time quantitative PCR

Transcript levels in WT and *clpc2-2* leaves were measured as described in Morcillo et al. (2024) using the primers listed in **Supplemental Table 8**. Comparative threshold values were normalized to *ACTIN2* internal control.

### Statistical analysis

Unless otherwise indicated, the data presented are the means (± SE) obtained from 3-4 independent experiments, with 3 replicates in each experiment. The significance of differences between the control and the transgenic lines was evaluated statistically with the Student’s t-test in the SPSS software. Differences were considered significant at a probability level of P<0.05.

## CONTRIBUTIONS

J P-R designed the experiments and analyzed the data; J L-L, AB, N DD, P T, E B-F, FJ M, G A, C E A-P, F I J-R, P P, D L, E C, I E, L L-S, A F-G, V C-R and RJL M performed most of the experiments and discussed the data; J L-L and J P-R wrote the article with contributions from all the authors; J P-R conceived the project and research plans.

## ACKNOWLEDGEMENTS

This work was supported by the Ministerio de Ciencia e Innovación y Universidades (MCIU) and Agencia Estatal de Investigación (AEI) / 10.13039/501100011033/ (grants PID2022-137292NB-I00 to J P-R and PID2023-150687NB-I00 to P.P) and by. J L-L acknowledges a pre-doctoral fellowship funded by MCIU/AEI/10.13039/501100011033 and by “ESF Investing in your future”

## SUPPLEMENTAL FIGURES

**Supplemental Figure 1:** (A) External phenotype of on soil-grown WT, *clpc1-1, clpc2-2* and three independent lines of *clpc2-2 promCLPC2:CLPC2* plants. (B) Time series for fresh weight (FW) of rosettes of WT, *clpc1-1, clpc2-2* and one representative line (line #1) of *clpc2-2 promCLPC2:CLPC2* plants. Values are means ± SE determined from three independent experiments using eight plants in each experiment.

**Supplemental Figure 2: Developmental characterization of *clpc2-2* seeds.** *2222*

**Supplemental Figure 3:** Relative abundance of *FTSHI4, PGDH2, PDE334, PETC, FD2, NADP-MDH* and *FSD1* transcripts as determined by RT-qPCR in leaves of WT and *clpc2-2* plants. Values are means ± SE for three biological replicates (each a pool of four plants) obtained from three independent experiments.

**Supplemental Figure 4: Stages in the construction of the *promCLPC2:CLPC2* and *35S:CLPC2* plasmids.** Primers used for PCR amplification of the 2.0 kbp immediately upstream of the translation start codon of CLPC2 and a complete *CLPC2* cDNA obtained from the RIKEN Arabidopsis cDNA collection (Seki et al., 1998, 2002) are listed in **Supplemental Table 7**.

**Supplemental Figure 5: *CLPC2* genotyping of WT, *clpc2-2, clpc2-2 promCLPC2:CLPC2* and *35S:CLPC2* plants.** (A) PCR analyses of WT, *clpc2-2, clpc2-2 promCLPC2:CLPC2* and *35S:CLPC2* plants. The following primer pairs (cf. **Supplemental Table 7**) were used for these analyses: clpc2seq2/FL84 (left gel), PJ02/8474 (central gel) and 35S_5/FL84 (right gel). (B) Schemes illustrating *CLPC2* structure in WT, *clpc2-2, clpc2-2 promCLPC2:CLPC2* and *35S:CLPC2* plants. Protein-coding exons are represented by black boxes. T-DNA insertion site in *clpc2-2* is indicated precisely, but size is not to scale.

## REFERENCES

Ameztoy, K., Baslam, M., Sánchez-López, Á.,… Pozueta-Romero, J. (2019) Plant responses to fungal volatiles involve global post-translational thiol redox proteome changes that affect photosynthesis. Plant Cell Environ. 42: 2627–2644

Ameztoy, K., Sánchez-López, Á.M., Muñoz, F.J., Bahaji, A., Almagro, G., Baroja-Fernández, E., Gámez-Arcas, S., De Diego, N., Doležal, K., Novák, O., Pěnčík, A., Alpízar, A., Rodríguez-Concepción, M., Pozueta-Romero, J. (2021) Proteostatic regulation of MEP and shikimate pathways by redox-activated photosynthesis signaling in plants exposed to small fungal volatiles. Front. Plant Sci. doi: 10.3389/fpls.2021.637976. https://doi.org/10.3389/fpls.2021.637976

Arsova, B., Hoa, U.,, Wimmelbacher, M., Greiner, E., Ústün, S., Melzer, M., Petersen, K., Lein, W., Börnke, F. (2010) Plastidial thioredoxin *z* interacts with two fructokinase-like proteins in a thiol-dependent manner: evidence for an essential role in chloroplast development in *Arabidopsis* and *Nitotiana benthamiana*. Plant Cell 22: 1498–1515

Bahaji, A., Almagro, G., Ezquer, I., Gámez-Arcas, S., Sánchez-López, Á.M., Muñoz, F.J., Barrio, R.J., Sampedro, M.C., De Diego, N., Spíchal, L., Doležal, K., Tarkowská, D., Caporali, E., Mendes, M.A., Baroja-Fernández, E., Pozueta-Romero, J. (2018) Plastidial phosphoglucose isomerase is an important determinant of seed yield through its involvement in gibberellin-mediated reproductive development and storage reserve biosynthesis in Arabidopsis. Plant Cell 30: 2082–2098.

Bali, S., Gautam, A., Dhiman, A., Michael, R., Dogra, V. (2023) Salicylate and jasmonate intertwine in ROS-triggered chloroplast-to-nucleus retrograde signaling. Physiol. Plantarum. 175(5):e14041. doi: 10.1111/ppl.14041

Baud, S., Dubreucq, B., Miquel, M., Rochat, C., and Lepiniec, L. (2008). Storage reserve accumulation in *Arabidopsis*: metabolic and developmental control of seed filling. Arabidopsis Book. doi:10.1199/tab.0113.

Baud, S., Wuillème, S., Dubreucq, B., de Almeida, A., Vuagnat, C., Lepiniec, L., Miquel, M., and Rochat, C. (2007) Function of plastidial pyruvate kinases in seeds of *Arabidopsis thaliana*. Plant J. 52: 405–419

Boeglin, M., Fuglsang, A.T., Luu, D-T., Sentenac, H., Gaillard, I., Chérel, I. (2016) Reduced expression of *AtNUP62* nucleoporin gene affects auxin response in *Arabidopsis*. BMC Plant Biol. 16:2 DOI 10.1186/s12870-015-0695-y

Chen, M., Choi, Y., Voytas, D.F., Rodermel, S. (2000) Mutations in the Arabidopsis *VAR2* locus cause leaf variegation due to the loss of a chloroplast FtsH protease. Plant J. 22: 303–313

Clough, S.J., Bent, A.F., (1998) Floral dip: A simplified method for Agrobacterium-mediated transformation of *Arabidopsis thaliana*. Plant J. 16: 735–743. 10.1046/j.1365-313X.1998.00343.x

Constan, D., Froehlich, J.E., Rangarajan, S., Keegstra, K. (2004) A stromal Hsp100 protein is required for normal chloroplast development and function in Arabidopsis. Plant Physiol. 136: 3605–3615

Cortleven, A., Valcke, R. (2012) Evaluation of the photosynthetic activity in transgenic tobacco plants with altered endogenous cytokinin content: lessons from cytokinin. Physiol. Plantarum 144: 394–408

Dvorák, P., Krasylenko, Y., Ovecka, M., Basheer, J., Zapletalová, V., Samaj, J., Takác, T. (2020) *In vivo* light-sheet microscopy resolves localization patterns of FSD1, a superoxide dismutase with function in root development and osmoprotection. Plant Cell Environ. 44: 68–87

Flexas, J., Díaz-Espejo, A., Conesa, M. A., Coopman, R. E., Douthe, C., Gago, J., Gallé, A., Galmés, J., Medrano, H., Ribas-Carbo, M., Tomàs, M., Niinemets, Ü. (2016). Mesophyll conductance to CO_2_ and Rubisco as targets for improving intrinsic water use efficiency in C3 plants. Plant Cell Environ. 39: 965–982

Flores-Pérez, U., Bédard, J., Tanabe, N., Lymperopoulos, P., Clarke, A.K., Jarvis, P. (2016) Functional analysis of the Hsp93/ClpC chaperone at the chloroplast envelope. Plant Physiol. 170: 147–162

Gámez-Arcas, S., Baroja-Fernández, E., García-Gómez, P., Muñoz, F.J., Almagro, G., Bahaji, A., Sánchez-López, A.M., Pozueta-Romero, J. (2022a) Action mechanisms of small microbial volatile compounds in plants. J. Exp. Bot. 73: 498–510. doi: 10.1093/jxb/erab463

Gámez-Arcas, S., Muñoz, F.J., Ricarte-Bermejo, A., Sánchez-López, A.M., Baslam, M., Baroja-Fernández, E., Bahaji, A., Almagro, G., De Diego, N., Dolezal, K., Novák, O., Leal-López, J., Mrocillo, R.J., Castillo, A.G., Pozueta-Romero, J. (2022b) Glucose-6-P/phosphate translocator2 mediates the phosphoglucose-isomerase-independent response to microbial volatiles. Plant Physiol. 190: 2137–2154

Gámez-Arcas, S., Muñoz, F.J., Serrato, A.J., Sánchez-López, Á.M., Baroja-Fernández, E., Goizeder Almagro, Abdellatif Bahaji, Leal-López, J., Morcillo, R.J.L., Pozueta-Romero, J. (2025) The microbial volatile-responsive, redox-sensitive Cys95 residue is an important structural and functional determinant of the Calvin-Benson fructose-1,6-bisphosphatase in Arabidopsis. Plant Cell Environ. 48: 6941–6951.

García-Gómez, P., Almagro, G., Sánchez-López, A.M., Bahaji, A., Ameztoy, K., Ricarte-Bermejo, A., Baslam, M., Antolín, M.C., Urdiain, A., López-Belchi, M.D., López-Gómez, P., Morán, J.F., Garrido, J., Muñoz, F.J., Baroja-Fernández, E., Pozueta-Romero, J. (2019) Volatile compounds other than CO_2_ emitted by different microorganisms promote distinct posttranscriptionally regulated responses in plants. Plant Cell Environ. 42: 1729–1746

García-Gómez, P., Bahaji, A., Gámez-Arcas, S., Muñoz, F.J., Sánchez-López, Á.M., Almagro, G., Baroja-Fernández, E., Ameztoy, K., De Diego, N., Ugena, L., Spíchal, L., Dolezal, K., Hajirezaei, M-R., Romero, L.C., García, I., Pozueta-Romero, J. (2020) Volatiles from the fungal phytopathogen *Penicillium aurantiogriseum* modulate root metabolism and architecture through proteome resetting. Plant Cell Environ. 43: 2551–2570 doi: 10.1111/pce.13817

Gonzalez-Guzmán, M., Pizzio, G.A., Antoni, R., Vera-Sirera, F., Merilo, E., Bassel, G.W., Fernández, M.A., Holdsworth, M.J., Perez-Amador, M.A., Kollist, H., Rodríguez, P.L. (2012) *Arabidopsis* PYR/PYL/RCAR receptors play a major role in quantitative regulation of stomatal aperture and transcriptional response to abscisic acid. Plant Cell 24: 2483–2496

Isono, E., Li, J., Pulido, P., Siao, W., Spoel, S.H., Wuang, Z., Zhuang, X., Trujillo, M. (2024) Protein degrons and degradation: Exploring substrate recognition and pathway selection in plants. Plant Cell 36: 3074–3098

Jasso-Robles, F.I., Aucique-Perez, C.E., Zelikovic, S.C., Saiz-Fernández, I., Klimes, P., De Diego, N. (2025) The loss-of-function of *AtNATA2* enhances *AtADC2-*dependent putrescine biosynthesis and priming, improving growth and salinity tolerance in Arabidopsis. Physiol. Plantarum doi: 176(6):e14603. doi: 10.1111/ppl.14603

Kieber, J.J., Schaller, G.E. (2014) Cytokinins. The Arabidopsis Book 12, e0168. 10.1199/tab.0168

Kikuchi, S., Asakura, Y., Imai, M., Nakahira, Y., Kotani, Y., Hashiguchi, Y., Nakai, Y., Takafuji, K., Bédard, J., Hirabayashi-Ishioka, Y., Mori, H., Shiina, T., Nakai, M. (2018) A Ycf2-FtsHi heteromeric AAA-ATPase complex is required for chloroplast protein import. Plant Cell 30: 2677–2703

Kim, J., Olinares P.D., Oh, S-h., Ghisaura, S., Poliakov, A., Ponnala, L., van Wijk, K.J. (2013) Modified clp protease complex in the ClpP3 null mutant and consequences of chloroplast development and function in Arabidopsis. Plant Physiol. 162: 157k–179

Kim, J., Rudella, A., Ramirez Rodríguez, V., Zybailov, B., Olinares., P.D.B, van Wijk, K.J. (2009) Subunits of the plastid ClpPR protease complex have differential contributions to embryogenesis, plastid biogenesis, and plant development in Arabidopsis. Plant Cell 21: 1669–1692

Kovacheva, S., Bédard, J., Wardle, A., Patel, R., Jarvis, P. (2007) Further *in vivo* studies on the role of the molecular chaperone, Hsp93, in plastid protein import. Plant J. 50: 364–379

Liao, J-Y. R., Friso, G. Forsythe, E.S., Michel, E.J.S., Williams, A.M., Boguraev, S.S., Ponnala, L., Sloan, D.B., van Wijk, K.J. (2022) Proteomics, phylogenetics, and coexpression analyses indicate novel interactions in the plastid CLP chaperone-protease system. J. Biol. Chem. 298: 101609

Liu, J., Wang, P., Liu, B., Feng, D., Zhang, J., Su, J., Zhang, Y., Wang, J-F., Wang, H-B. (2013) A defeciency in chloroplastic ferredoxin 2 facilitates effective photosynthetic capacity during long-term high light acclimation in *Arabidopsis thaliana*. Plant J. 76: 861–874

Loperfido, D., Zumajo-Cardona, C., Commisso, M., Guzzo, F., Ambrose, B., Colombol, L., Ezquer, I. (2025) The role of cytokinin receptors in Arabidopsis thaliana seed development and how they affect the metabolomoc profile. Planta 262(3):59. doi: 10.1007/s00425-025-04745-7.

López, B., Izquierdo, Y., Cascón, T., Zamarreño, a.M., García-Mina,. M., Pulido, P., Castresana, C. (2024) Mutant *noxy8* exposes functional specificities between the chloroplast chaperones CLPC1 and CLPC2 in the response to organelle stress and plant defence. Plant Cell Environ. 47: 2336–2350. doi: 10.1111/pce.14882

Lu, X., Zhang, D., Li, S., Su, Y., Liang, Q., Meng, H., Shen, S., Fan, Y., Liu, C., Zhang C. (2014) FtsHi4 is essential for embryogenesis due to its influence on chloroplast developmentin Arabidopsis. PLOS One 15(2): e0229232. 10.1371/journal.pone.0229232

Maiwald, D., Dietzmann, A., Jahns, P., Pesaresi, P., Joliot, P, Joliot, A., Levin, J.Z., Salamini, F., Leister, D. (2003) Knock-out of the genes coding for the rieske protein and the ATP-synthase g-subunit of Arabidopsis. Effects on photosynthesis, thylakoid protein composition and nuclear chloroplast gene expression. Plant Physiol. 133: 191–202

Mateo, A., Funck, D., Mühlenbock, P., Kular, B., Mullineaux, P.M., Karpinski, S. (2006) Controlled levels of salicylic acid are required for optimal photosynthesis and redox homeostasis. J. Exp. Bot. 57: 1795–1807

Meinke, D (2019) Genome-wide identification of *EMBRYO-DEFECTIVE (EMB)* genes required for growth and development in Arabidopsis. New Phytol. 226: 306–325

Meng, L., Wong, J.H., Feldman, L.J., Lemaux, P.G., Buchanan, B.B. (2010) A membrane-associated thioredoxin required for plant growth moves from cell to cell, suggestive a role in intercellular communication. Proc. Natl. Acad. Sci. USA 107: 3900–3905

Montandon, C., Friso, G., Liao, J.Y.R., Choi, J., van Wijk, K.J. (2019) In vivo trapping of proteins interacting with the chloroplast CLPC1 chaperone: potential substrates and adaptors. J. Proteome Res. 18: 2585–2600

Morcillo, R.J.L., Leal-López, J., Férez-Gómez, A., López-Serrano, L., Baroja-Fernández, E., Gámez-Arcas, S., Tortosa, G., López, L.E., Estévez, J.M., Doblas, V.G., Frías-España, L., García-Pedrajas, M.D., Sarmiento-Villamil, J., Pozueta-Romero, J. (2024) RAPID ALKALINIZATION FACTOR 22 is a key modulator of the root hair growth responses to fungal ethylene emissions in Arabidopsis. Plant Physiol. 196: 2890–2904

Nishimura, K., Asakura, Y., Friso, G., Kim, J., Oh, S-h., Rutschow, H., Ponnala, L., van Wijk, K.J. (2013) ClpS1 is a conserved substrate selector for the chloroplast Clp protease system in *Arabidopsis*. Plant Cell 25: 2276–2301

Nishimura, K., van Wijk, K.J. (2015) Organization, function and substrates of the essential Clp protease system in plastids. Biochim. Biophys. Acta 1847: 915–930

Park, S., Rodermel, S.R. (2004) Mutations in ClpC2/Hsp100 suppress the requirement for FtsH in thylakoid membrane biogenesis. Proc. Natl. Acad. Sci. USA 101: 12765–12770

Parry, G. (2014) Components of the *Arabidopsis* nuclear pore complex play multiple diverse roles in control of plant growth. J. Exp. Bot. 65: 6057–6067

Peltier, J.B., Ripoll, D.R., Friso, G., Rudella, A., Cai, Y., Ytterberg, J., Giacomelli, L., Pillardy, J., Van Wijk, K.J. (2004) Clp protease complexes from photosynthetic and non-photosynthetic plastids and mitochondria of plants, their predicted three-dimensional structures, and functional implications. J. Biol. Chem. 279: 4768–4781.

Pulido, P., Llamas, E., Llorente, B., Ventura, S., Wright, L.P., Rodriguez-Concepcion, M. (2016) Specific Hsp100 Chaperones determine the fate of the first enzyme of the plastidial isoprenoid pathway for either refolding or degradation by the stromal Clp protease in Arabidopsis. PLOS Genet. 12(1): e1005824

Rojo, E., Gillmor, CS., Kovaleva, V., Somerville, C.R., Raikhel, N.V. (2001) *VACUOLELESS1* is an eseential gene required for vacuole formation and morphogenesis in *Arabidopsis*. Developmental Cell 1: 303–310

Sánchez-López, Á.M., Bahaji, A., De Diego, N., Baslam, M., Li, J., Muñoz, F.J., Almagro, G., García-Gómez, P., Ameztoy, K., Ricarte-Bermejo, A., Novák, O., Humplík, J.F., Spíchal, L., Doležal, K., Ciordia, S., Mena, M.C., Navajas, R., Baroja-Fernández, E., Pozueta-Romero, J. (2016a) Arabidopsis responds to *Alternaria alternata* volatiles by triggering plastid phosphoglucose isomerase-independent mechanisms. Plant Physiol. 172: 1989–2001.

Sánchez-López, Á.M., Baslam, M., De Diego, N., et al. (2016b) Volatile compounds emitted by diverse phytopathogenic microorganisms promote plant growth and flowering through cytokinin action. Plant, Cell Environ. 39: 2592–2608

Savchenko, T., Kolla, V.A., Wang, C-Q., Nasafi, Z., Hicks, D.R., Phandungchob, B., Chehab, W.E., Brandizzi, F., Froehlich, J., Dehesh, K. (2014) Functional convergence of oxylipin and abscisic acid pathways controls stomatal closure in response to drought. Plant Physiol. 164: 1151–1160

Schreier, T., B.,, Cléry, A., Schläfli, M., Galbier, F., Stadler, M., Demarsy, E., Albertine, D., Maier, B.A., Kessler, F., Hörtsensteiner, S., Zeeman, S.C., Kötting, O. (2018) Plastidial NAD-dependent malate dehydrogenase: a moonlighting protein involved in early chloroplast development through its interaction with an FtsH12-FtsHi protease complex. Plant Cell 30: 1745–1769

Seki, M., Carninci, P., Nishiyama, Y., Hayashizaki, Y., Shinozaki, K. (1998) High-efficiency cloning of Arabidopsis full-length cDNA by biotinylated CAP trapper. Plant J 15: 707–720

Seki, M., Narusaka, M., Kamiya, A., Ishida, J., Satou, M., Sakurai, T., Nakajima, M., Enju, A., Akiyama, K., Oono, Y., et al. (2002) Functional annotation of a full-length Arabidopsis cDNA collection. Science 296: 141–145

Sjögren, L.L.E., MacDonald, T.M., Sutinen, S., Clarke, A.K. (2004) Inactivation of the *CLPC1* gene encoding a chloroplast Hsp100 molecular chaperone causes growth retardation, leaf chlorosis, lower photosynthetic activity and a specific reduction in photosystem content. Plant Physiol. 136: 4114–4126

von Caemmerer, S., Farquhar, G.D. (1981) Some relationships between the biochemistry and photosynthesis and the gas exchange of leaves. Planta 153: 376–387

Vrobel, O., Zelikovic, S.C., Dehner, J., Spíchal, L., De Diego, N., Tarkowski, P. (2024) Multi-class plant hormone HILIC-MS/MS analysis coupled with high-throughput phenotyping to investigate plant-environment interactions. Plant J. doi: 10.1111/tpj.17010

Welsch, R., Zhou, X., Yuan, H., Álvarez, D., Sun, T., Schlossarek, D., Yang, Y., Shen, G., Zhang, H., Rodríguez-Concepción, M., Thannhauser, T.W., Li, L. (2018) Clp protease and OR directly control the proteostasis of phytoene synthase, the crucail enzyme for carotenoid biosynthesis in *Arabidopsis*. Mol. Plant 11: 149–162

Yokochi, Y., Yoshida, K., Hahn, F., Miyagi, A., Wakabayashi, K., Kawai-Yamada, M., Weber, A.P.M., Hisabori, T. (2021) Redox regulation of NADP-malate dehydrogenase is vital for land plants under fluctuating light environment. Proc. Natl. Acade. Sci. USA 118(6):e2016903118. doi: 10.1073/pnas.2016903118

Yu, F., Liu, X., Aksheikh, M., Park, S., Rodermel, S. (2008) Mutations in *SUPPRESSOR OF VARIEGATION1*, a factor required for normal chloroplast translation, suppress *var2-*mediated leaf variegation in *Arabidopsis*. Plant Cell 20: 1786–1804

Zaltsman, A., Ori, N., Adam, Z. (2005) Two types of FtsH protease subunits are required for chloroplast biogenesis and photosystem II repair in Arabidopsis. Plant Cell 17: 2782–2790

Zybailov, B., Friso, G., Kim, J., Rudella, A., Ramírez Rodríguez, V., Asakura, Y., Sun, Q., van Wijk, K.J. (2009) Large scale comparative proteomics of a chloroplast Clp protease mutant reveals folding stress, altered protein homeostasis, and feedback regulation of metabolism. Mol. Cell Proteomics. 8: 1789–1810

