## Supplemental Figure 1 for "CLPC2 plays specific roles in CLP complex-mediated regulation of growth, photosynthesis, embryogenesis and response to growth-promoting microbial compounds"

A

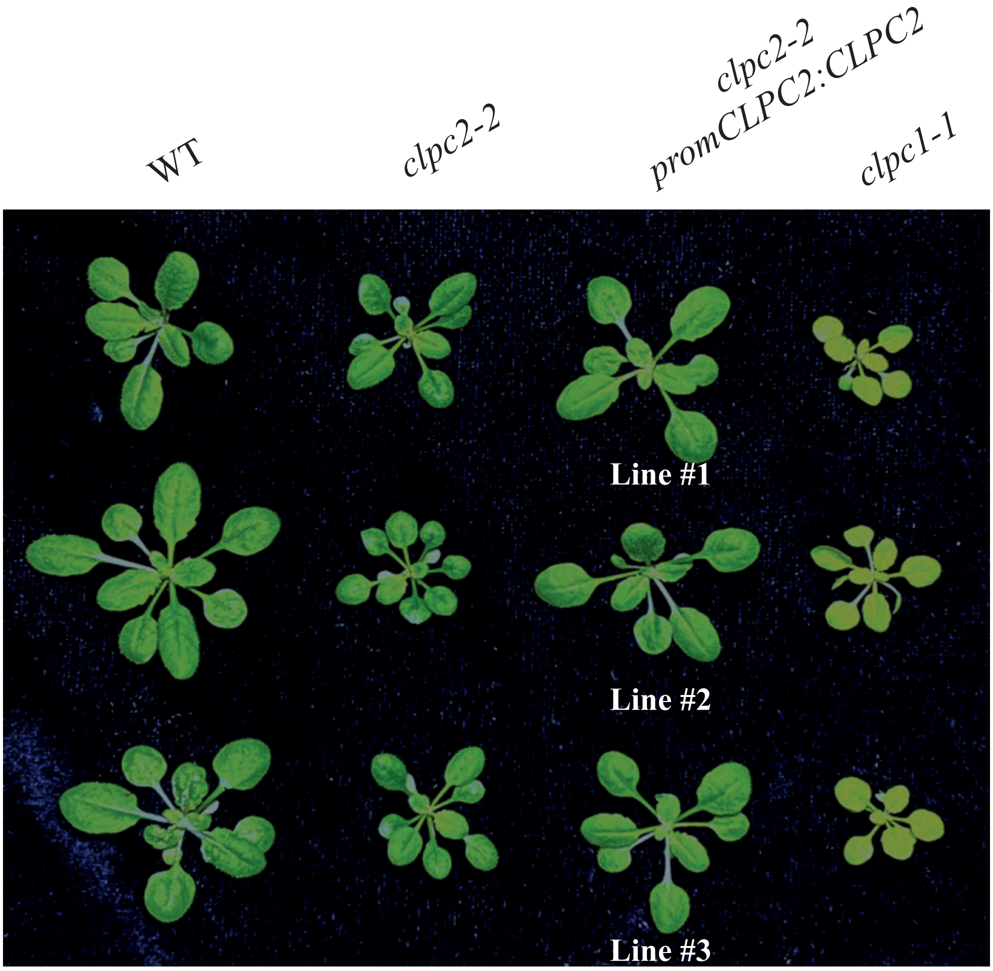


B

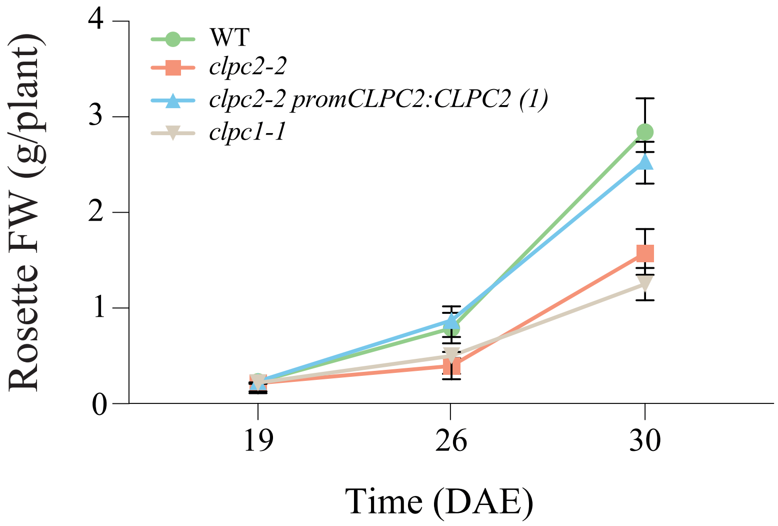


Supplemental Figure 1: (A) External phenotype of on soil-grown WT, *clpc1-1*, *clpc2-2* and three independent lines of *clpc2-2 promCLPC2:CLPC2* plants. (B) Time series for fresh weight (FW) of rosettes of WT, *clpc1-1, clpc2-2* and one representative line (line #1) of *clpc2-2 promCLPC2:CLPC2* plants. Values are means ± SE determined from three independent experiments using eight plants in each experiment.
