## Supplemental Figure 2 for "CLPC2 plays specific roles in CLP complex-mediated regulation of growth, photosynthesis, embryogenesis and response to growth-promoting microbial compounds"

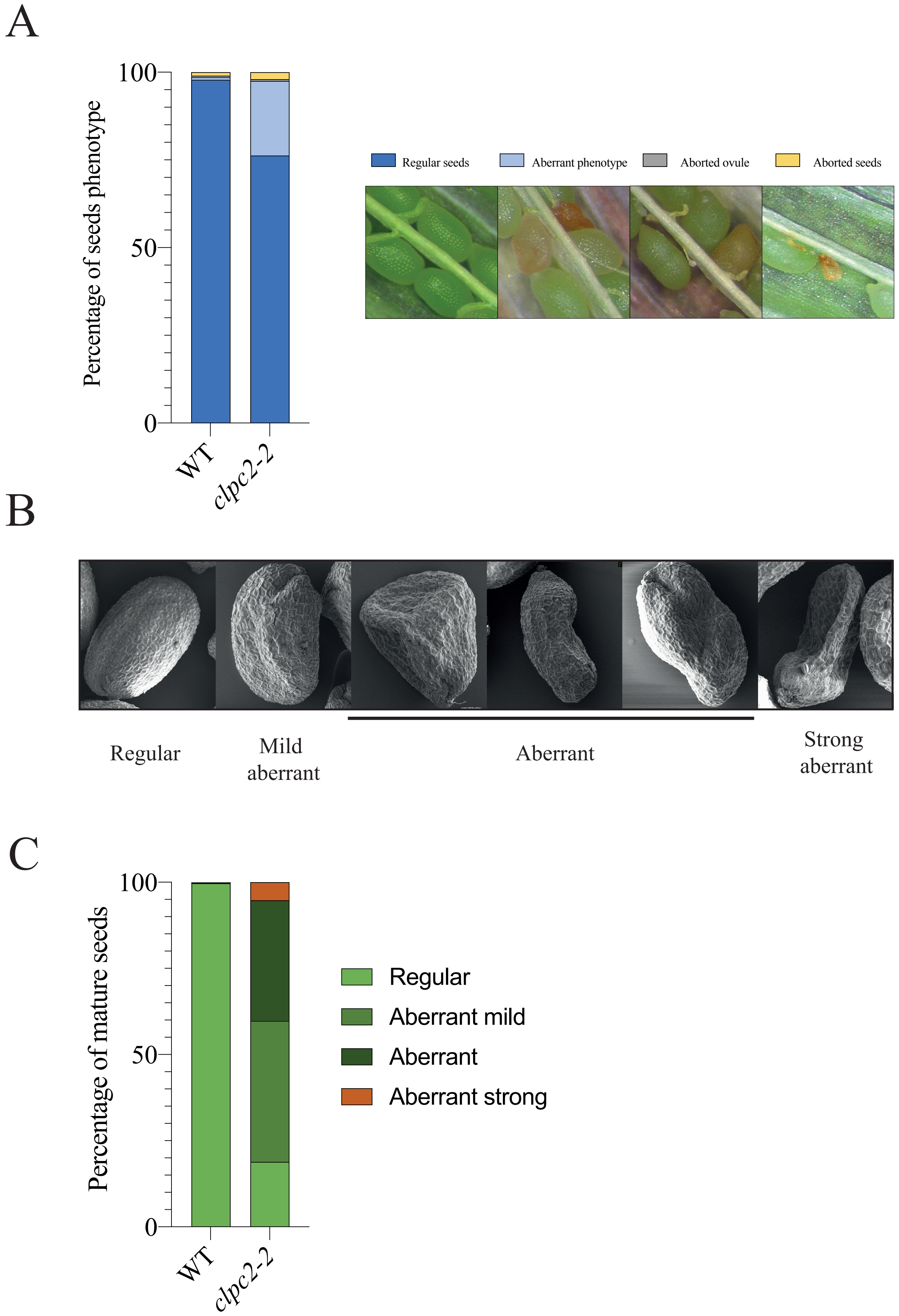


Supplemental Figure 2: Developmental characterization of *clpc2-2* seeds. (A) Representative siliques from WT and *clpc2-2* plants at 15 days after pollination (DAP), along with the percentage of seed developmental defects. A total of 22 siliques per genotype were analyzed, derived from 5–6 independent plants. The proportion of “normal” and “aberrant” seed phenotypes shows statistically significant differences between WT and *clpc2-2* mutants (Oneway ANOVA followed by Tukey HSD test was used for analyzing significance), whereas the percentage of aborted ovules and seeds does not differ significantly between genotypes. (B) Scanning electron microscopy analysis of seed morphology (C) Quantification of the different types of aberrant phenotypes of WT and *clpc2-2* seeds. Approximately 800 seeds per genotype were analyzed, derived from 5 independent plants. All reported categories show statistically significant differences between WT and *clpc2-2* mutants (Oneway ANOVA followed by Tukey HSD test was used for analyzing significance).
