## Supplemental Figure 3 for "CLPC2 plays specific roles in CLP complex-mediated regulation of growth, photosynthesis, embryogenesis and response to growth-promoting microbial compounds"

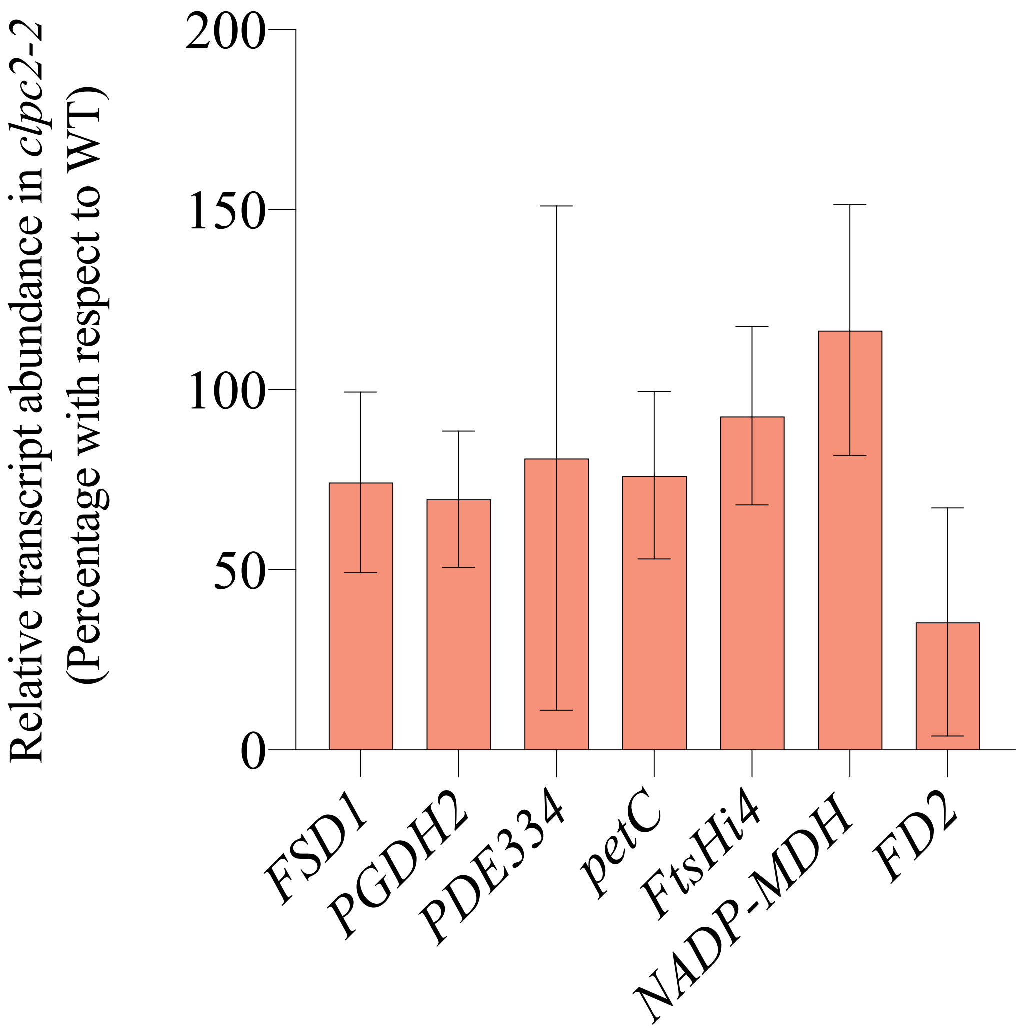


Supplemental Figure 3: Relative abundance of *FTSHI4, PGDH2, PDE334, PETC, FD2, NADP-MDH* and *FSD1* transcripts as determined by RT-qPCR in leaves of WT and *clpc2-2* plants. Values are means ± SE for three biological replicates (each a pool of four plants) obtained from three independent experiments.
