## Supplemental Figure 4 for "CLPC2 plays specific roles in CLP complex-mediated regulation of growth, photosynthesis, embryogenesis and response to growth-promoting microbial compounds"

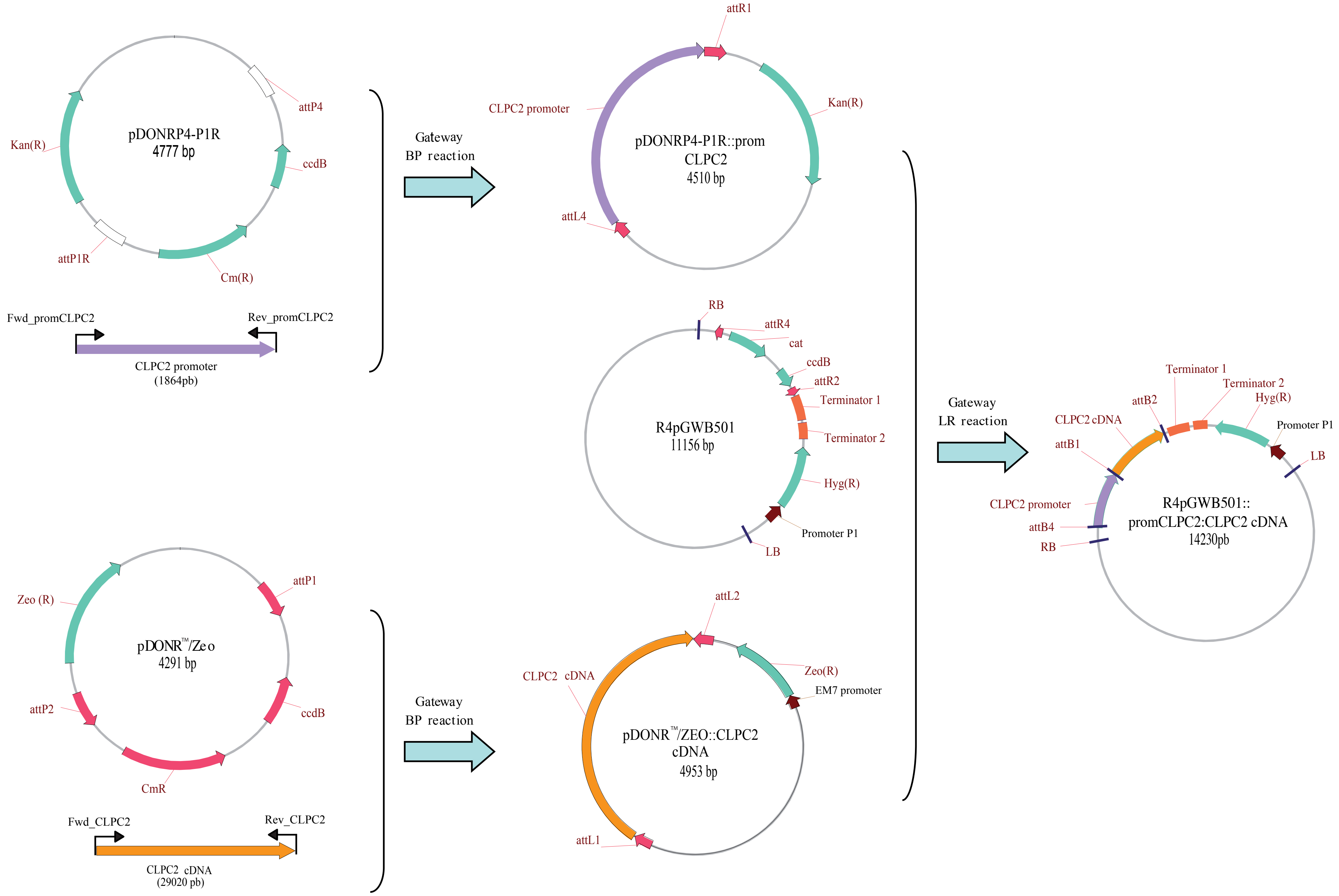


Supplemental Figure 4

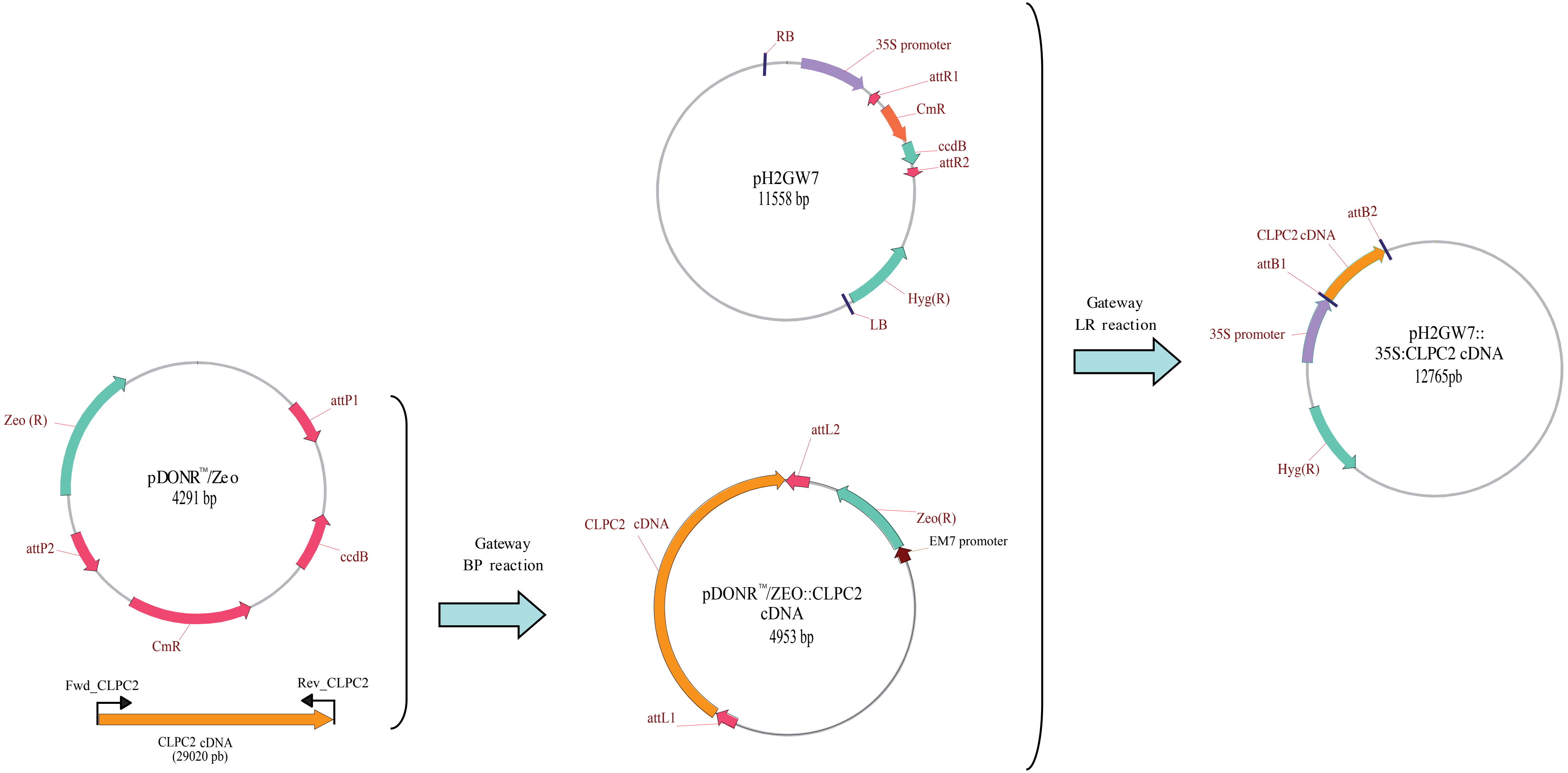


Supplemental Figure 4: Stages in the construction of the *promCLPC2:CLPC2* and *35S:CLPC2* plasmids. Primers used for PCR amplification of the 2.0 kbp immediately upstream of the translation start codon of CLPC2 and a complete CLPC2 cDNA obtained from the RIKEN Arabidop-
