## Supplemental Figure 5 for "CLPC2 plays specific roles in CLP complex-mediated regulation of growth, photosynthesis, embryogenesis and response to growth-promoting microbial compounds"

A

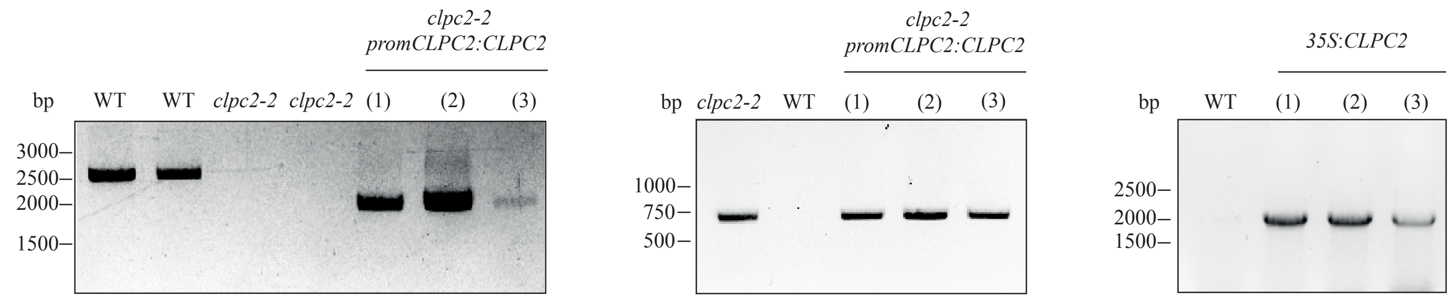


B

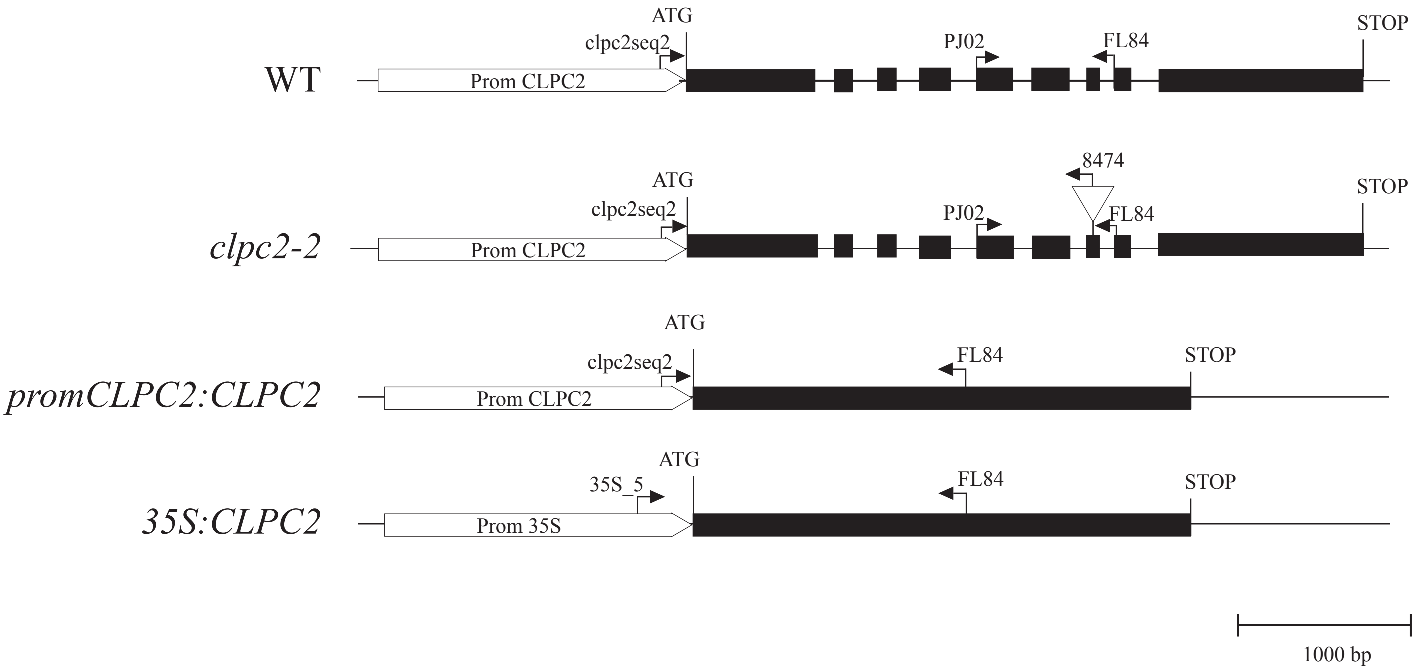


Supplemental Figure 5: CLPC2 genotyping of WT, *clpc2-2, clpc2-2 promCLPC2:CLPC2* and *35S:CLPC2* plants. (A) PCR analyses of WT, *clpc2-2, clpc2-2 promCLPC2:CLPC2* and *35S:CLPC2* plants. The following primer pairs (cf. Supplemental Table 7) were used for these analyses: clpc2seq2/FL84 (left gel), PJ02/8474 (central gel) and 35S_5/FL84 (right gel). (B) Schemes illustrating CLPC2 structure in WT, *clpc2-2, clpc2-2 promCLPC2:CLPC2* and *35S:CLPC2*
